# Untargeted Metabolomics Reveals Organ-Specific and Extraction-Dependent Metabolite Profiles in Endemic Tajik Species *Ferula violacea* Korovin

**DOI:** 10.1101/2025.08.24.671964

**Authors:** Sylhiya Mavlonazarova, Kenneth Acosta, Rinat Abzalimov, Saidbeg Satorov, Vyacheslav Dushenkov

## Abstract

*Ferula violacea* Korovin, an endemic Tajikistani plant with purported medicinal properties, remains understudied. This study employs untargeted metabolomics to characterize the metabolite profiles of ethanol extracts and juices from *F. violacea* roots and seeds. In total, 540 distinct metabolites are putatively identified, 419 of which are previously unreported in the *Ferula* genus, representing a substantial expansion of its known chemical diversity. The most abundant metabolites are terpenoids, amino acid derivatives, and alkaloids. A particularly abundant group of daucane sesquiterpenoids, sharing a common (6-methyl-azulen-4-yl)cyclohexanecarboxylate substructure, is identified, including known metabolites such as ferutidin and ferutinin. Comparative analysis reveals organ-specific metabolic specialization: roots are enriched in terpenoids, whereas seeds exhibit higher concentrations of alkaloids and amino acids. Additionally, processing methods influence metabolite composition, with ethanol extracts being rich in terpenoids and amino acids, and juices displaying a greater diversity of phenylpropanoid-derived compounds. These findings expand the phytochemical richness of *F. violacea* and suggest its potential as a valuable source of bioactive compounds for pharmacological exploration.

**Significance statement:** This study presents the first comprehensive untargeted metabolomic analysis of *Ferula violacea*, identifying 419 previously unreported metabolites for the genus and significantly expanding its known chemical diversity. The findings reveal organ- and method-specific metabolic specialization, underscoring the plant’s potential as a valuable source of novel bioactive compounds for pharmacological exploration.

## 1. Introduction

With an estimated total plant chemical space encompassing up to 25.7 million metabolites (Hart, Gadiya et al. 2025), medicinal plants represent a rich repository of bioactive phytochemicals. These compounds remain pivotal in managing chronic diseases and addressing emerging infectious threats, such as COVID-19 (Dushenkov and Dushenkov 2022, Bogoyavlenskiy, Alexyuk et al. 2024). Their role in promoting long-term health and alleviating chronic conditions, including cancer, cardiovascular disease, and diabetes, is increasingly recognized (Ahmad and Karmakar 2023).

Tajikistan, located within the Mountains of Central Asia, is notable for its exceptional floristic diversity, hosting over 4,300 vascular plant species, including more than 1,400 endemics (Nowak, Świerszcz et al. 2020). Among these, the genus *Ferula* is one of the largest within the Apiaceae family, comprising approximately 227 species worldwide, primarily located in Central Asia (Safarov, Hisoriev et al. 2019, WFO 2024). Despite this diversity, scientific literature has focused predominantly on *F. tadshikorum* Pimenov, with limited attention to other endemic species such as *F. violacea* Korovin, *F. karategina* Lipsky ex Korovin, *F. koso-poljanskyi* Korovin, *F. linczevskii* Korovin, *F. decurrens* Korovin, and *F. botschantzevii* Korovin. This underrepresentation highlights a critical gap in the phytochemical and pharmacognostic characterization of Tajik *Ferula* species.

*Ferula violacea* Korovin, an endemic species confined to Tajikistan (Pimenov 2020), belongs to the genus, which includes medicinally important taxa such as *F. assa-foetida* L. known for producing asafoetida, a resinous exudate, with diverse biological activities, including antifungal, antiparasitic, antitumor, anti-inflammatory properties, and has been shown to induce apoptosis in colorectal cancer cell lines (Elarabany, Hamad et al. 2023). Consequently, continued phytochemical exploration of *Ferula* species is vital to unlocking their full therapeutic potential (Karimi, Jariani et al. 2024). Phytochemically, *Ferula* species are characterized by the presence of sulfur-containing compounds, monoterpenes, and sesquiterpenes. Sulfur compounds, prevalent in asafoetida-yielding species, exhibit antimicrobial, antifungal, and carminative effects, and are valued for treating respiratory and gastrointestinal ailments. Terpenoids such as α-pinene, β-eudesmol, limonene, and myrcene contribute to the genus’s aromatic profile and possess notable bioactivities, including antimicrobial, insecticidal, and antifungal effects. Only two constituents—umbelliprenin and galbanic acid—had previously been identified in acetone extracts of *F. violacea* (Kir’yanova, Sklyar et al. 1979). Reflecting the growing interest in the species’ phytochemistry, the volatile metabolites of its essential oil were recently analyzed using GC-MS (Ghaforzoda, Sharopov et al. 2025).

In the present study, we employed untargeted metabolomic profiling using UHPLC-QTOF mass spectrometry to elucidate the chemical composition of *F. violacea* roots and seeds. The objectives were to expand the current knowledge of *F. violacea* metabolic diversity, evaluate organ-specific metabolite distribution, and assess the influence of extraction methods on phytochemical yield. The findings of this study aim to establish a foundation for future pharmacological research and underscore the potential of *F. violacea* as a source of bioactive compounds.

## 2. Results

### 2.1. Identification of *F. violacea* Metabolites Using Untargeted Metabolomics

An untargeted metabolomic analysis of *F. violacea* was conducted using a Bruker maXis-II UHR-ESI-QqTOF mass spectrometer coupled with a Thermo Scientific Ultimate 3000 UHPLC system. A mass feature table was generated with Bruker MetaboScape software, and after data harmonization, subsequent processing and curation, 540 mass features remained (**Table S1**). Putative metabolites (Level 2a and Level 3 confidence, (Schymanski, Jeon et al. 2014, Schrimpe-Rutledge, Codreanu et al. 2016)) were identified by matching mass feature spectra to known compounds using both experimental and *in-silico* spectral libraries.

To ensure accurate feature identifications, assigned chemical structures were evaluated to confirm they represented biological metabolites rather than synthetic compounds. NP-likeness scores were calculated for assigned structures to assess their structural similarity to natural products; most structures (n = 470) scored above 0 (**Figure S1A**). To further verify their biological relevance, assigned structures were cross-referenced with the ChEBI database for primary metabolites and several plant natural product collections from the COCONUT database for plant secondary metabolites. Of the 540 assigned structures, only 3 were not found in any of the biochemical databases examined (**Figure S1B**). Collectively, these results support the accuracy of the assigned feature identifications as natural metabolites.

Newly reported metabolites were determined by cross-referencing public database records for previously identified natural products within the *Ferula* genus. *Ferula* metabolites were further classified according to their biosynthetic pathways, superclasses, and chemical classes (Table S1).

### 2.2. Chemical Diversity of *F. violacea*

#### 2.2.1. Chemical Diversity of F. violacea Metabolites

**Table 1** presents a comparative analysis of previously reported metabolites in *Ferula* species and those newly reported in this study. Our results reveal widening of the metabolite diversity across multiple biosynthetic pathways (Table 1). Terpenoids remained the most diverse metabolites, with 213 identified in *F. violacea*, of which 143 are newly reported. Similarly, shikimates and phenylpropanoids represent a major pathway, comprising 121 identified in this study, including 83 previously unreported structures. Notably, a pronounced increase was observed in alkaloids and their hybrids, with 56 newly reported metabolites. Likewise, the diversity of amino acids and peptide derivatives demonstrated marked enrichment from just 6 previously reported metabolites to 45 newly reported structures, all of which are novel for the genus. Additionally, nearly all identified polyketide-derived compounds (38) had not been previously reported in the *Ferula* genus (37). Other biosynthetic pathways, including those associated with fatty acids and carbohydrates, also exhibited growth in chemical diversity. Additionally, nearly all identified polyketide-derived compounds (38) had not been previously reported in the *Ferula* genus (37). Other biosynthetic pathways, including those associated with fatty acids and carbohydrates, also exhibited growth in chemical diversity.

**Table 1.**
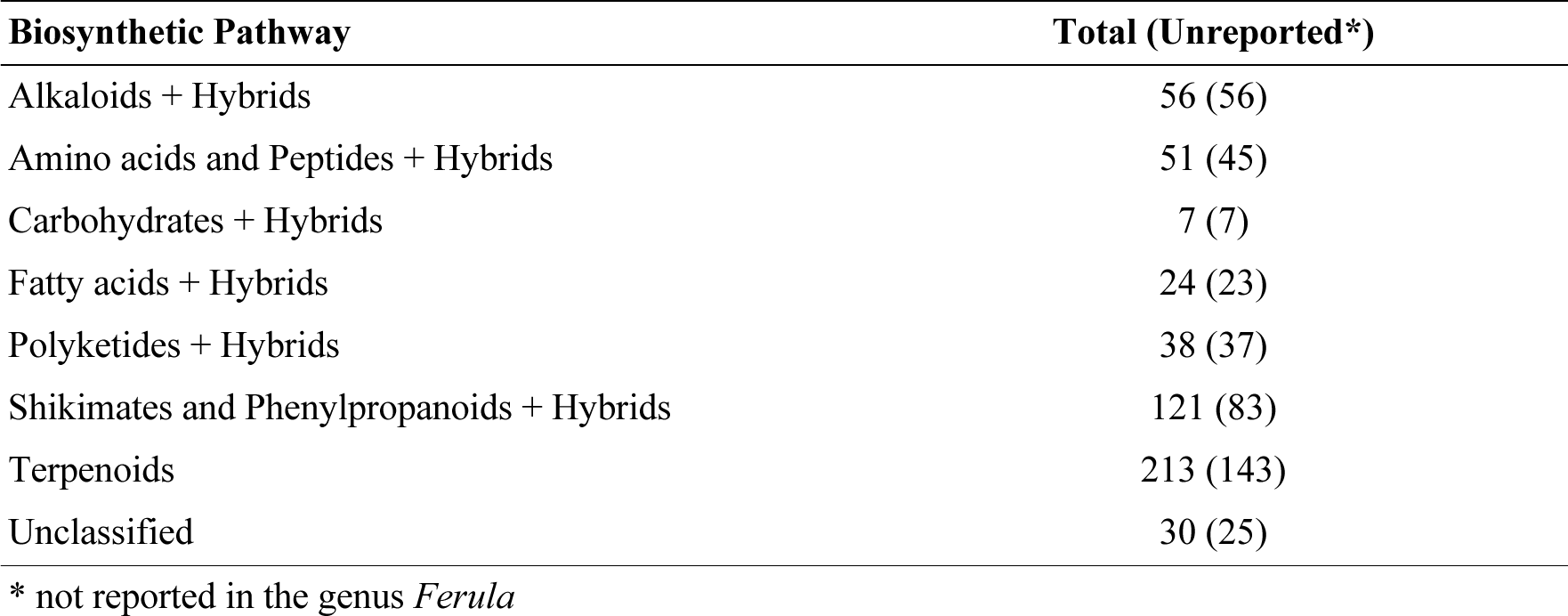
Number of metabolites from various natural product biosynthetic pathways in *F. violacea*.

**Table 2** presents the distribution of terpenoid superclasses within *F. violacea*, comparing previously reported terpenoids in *Ferula* with those newly reported in this study. Our findings indicate a substantial rise in terpenoid diversity, particularly within the sesquiterpenoid and monoterpenoid superclasses. Sesquiterpenoids and their hybrids remained the dominant superclass, with 115 identified in *F. violacea*, including 67 previously unreported metabolites. Monoterpenoids also exhibited considerable diversity, with 34 identified in this study, of which 21 are newly reported within the *Ferula* genus. Apocarotenoid diversity increased, with 1 previously reported in *Ferula* and 8 newly reported in this study. Diterpenoids (4 previously reported in *Ferula*, 12 unreported) and meroterpenoids (3 previously reported, 15 newly reported) demonstrated moderate increases in diversity. The steroid superclass showed a pronounced increase, with 10 identified steroids, almost all of which were previously unreported for *Ferula* (9). Triterpenoid diversity, however, remained relatively limited, with only 1 newly reported in this study, as well as carotenoids (C40) with 2 newly reported in this study.

**Table 2.**
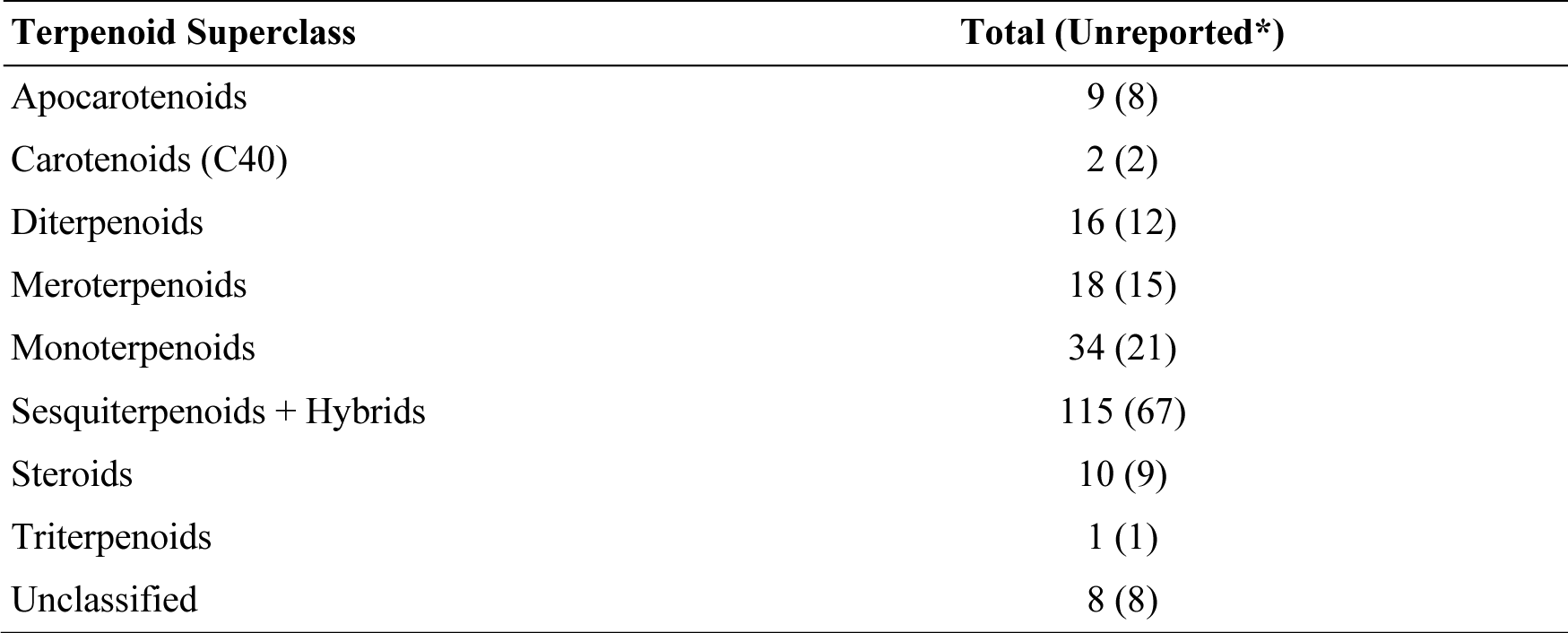
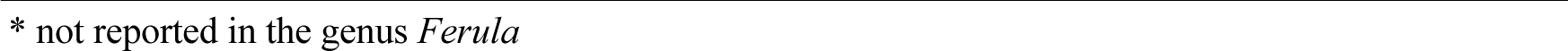
Number of metabolites from various terpenoid superclasses in *F. violacea*.

| Terpenoid Superclass | Total (Unreported*) |
| --- | --- |
| Apocarotenoids | 9 (8) |
| Carotenoids (C40) | 2 (2) |
| Diterpenoids | 16 (12) |
| Meroterpenoids | 18 (15) |
| Monoterpenoids | 34 (21) |
| Sesquiterpenoids + Hybrids | 115 (67) |
| Steroids | 10 (9) |
| Triterpenoids | 1 (1) |
| Unclassified | 8 (8) |

\* not reported in the genus *Ferula*

The distribution of *F. violacea* natural products derived from the shikimate and phenylpropanoid pathways is presented in **Table 3**, which compares previously reported metabolites in *Ferula* with those newly reported in this study. A significant number of previously unreported metabolites were identified from several superclasses, particularly phenolic acids, phenylpropanoids, and flavonoids. Phenolic acids (C6-C1) and their hybrids, known for their roles in antioxidant activity and signaling functions, demonstrated the most substantial increase in diversity, with 28 identified in this study, 21 of which had not been previously reported for *Ferula*. Similarly, phenylpropanoids (C6-C3) and their hybrids, which serve as key intermediates in lignin biosynthesis and the production of bioactive secondary metabolites, exhibited novel diversity, with 28 identified compounds, 17 of which are previously unreported within the *Ferula* genus. Coumarins showed a moderate increase in diversity, with 21 identified in *F. violacea*, including 9 newly reported metabolites. Flavonoids, known for their antioxidant and anti-inflammatory properties, have been previously reported in *Ferula*, with 16 identified in this study, 11 of which were newly reported. Lignans have been reported in *Ferula*, with 5 identified in this study, 4 of which are newly reported. Beyond these dominant superclasses, several previously unreported small molecule phenylpropanoid derivatives were identified in *F. violacea*, including 1 phenanthrenoids, 1 stilbenoid, and 1 styrylpyrone.

**Table 3.** Number of metabolites from various shikimate and phenylpropanoid superclasses in *F. violacea*.

| Shikimate and Phenylpropanoid Superclass | Total (Unreported*) |
| --- | --- |
| Coumarins + Hybrids | 21 (9) |
| Flavonoids | 16 (11) |
| Lignans | 5 (4) |
| Phenanthrenoids | 1 (1) |
| Phenolics acids (C6-C1) + Hybrids | 28 (21) |
| Phenylpropanoids (C6-C3) | 28 (17) |
| Small peptides | 1 (1) |
| Stilbenoids | 1 (1) |
| Styrylpyrones | 1 (1) |
| Unclassified | 19 (17) |

\* not reported in the genus *Ferula*

Alkaloids constitute a structurally diverse class of nitrogen-containing secondary metabolites with significant pharmacological potential. **Table 4** compares alkaloids identified in this study with those previously reported within the *Ferula* genus. Results reveal all 56 alkaloids identified in this study are newly reported to the *Ferula* genus. This includes 18 tryptophan-derived, 7 anthranilic acid-derived, 5 nicotinic acid-derived, and 3 tyrosine-derived, emphasizing the role of aromatic amino acid metabolism in *Ferula* alkaloid biosynthesis. Additionally, the identification of newly reported peptide alkaloids (1) and tetramate/peptide alkaloids (2), suggests previously unrecognized non-ribosomal peptide synthetase activity in *Ferula*. Beyond alkaloids derived from aromatic amino acids, our study identified ornithine- and lysine-derived alkaloids, potentially indicating an expanded biosynthetic capacity for piperidine and quinolizidine alkaloids in *Ferula*. Lysine-derived alkaloids and their hybrids increased to include 5 newly reported metabolites, while the presence of an ornithine-derived alkaloid suggests a possible link to polyamine metabolism, which is often implicated in stress responses and secondary metabolite regulation. A notable finding was the identification of 7 newly reported pseudoalkaloids, which likely originate from terpenoid, polyketide, or steroidal intermediates rather than classical amino acid biosynthesis.

**Table 4.** Number of metabolites from various alkaloid superclasses in *F. violacea*.

| Alkaloid Superclass | Total (Unreported*) |
| --- | --- |
| Anthranilic acid alkaloids | 7 (7) |
| Lysine alkaloids + Hybrids | 5 (5) |
| Nicotinic acid + Hybrids | 5 (5) |
| Ornithine alkaloids | 1 (1) |
| Peptide alkaloids | 1 (1) |
| Pseudoalkaloids | 7 (7) |
| Tetramate alkaloids/Peptide alkaloids | 2 (2) |
| Tryptophan alkaloids + Hybrids | 18 (18) |
| Tyrosine alkaloids + Hybrids | 3 (3) |
| Unclassified | 7 (7) |

\* not reported in the genus *Ferula*

#### 2.2.2. Metabolomic Profiling of F. violacea Identifies Abundant Terpenoids

The chemical composition of *F. violacea* was investigated by determining the relative abundance of various biosynthetic pathways and their associated superclasses. **Figure 1** shows the abundance of natural product biosynthetic pathways within *F. violacea* (**Figure 1A**), highlighting terpenoids, amino acids, and alkaloids as the most abundant metabolites recovered in this species.

**Figure 1.**
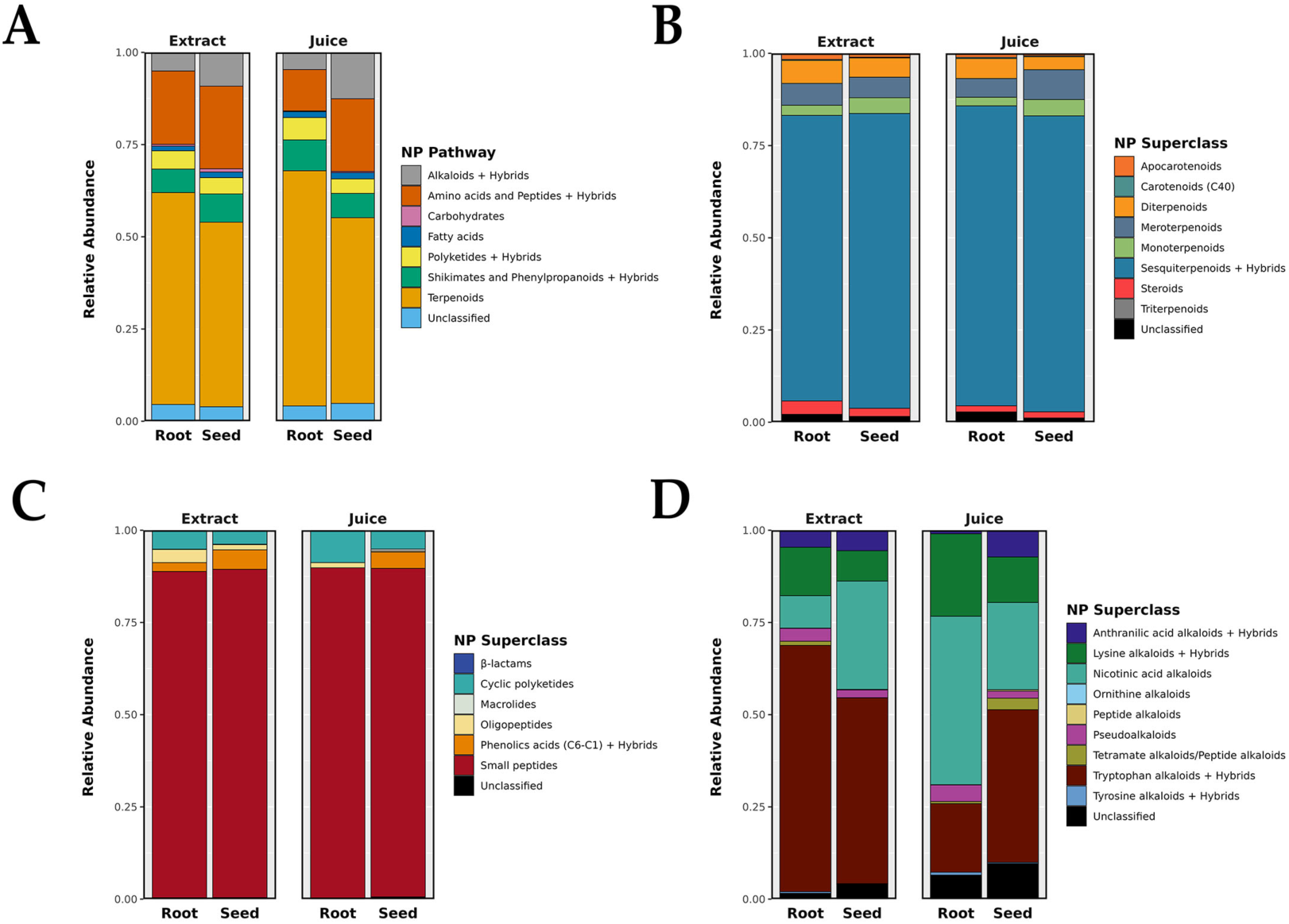
Metabolite Profiles of *F. violacea* Samples. (A) Relative abundance of natural product biosynthetic pathways. The intensities of metabolites within each biosynthetic pathway were grouped and summed. Metabolites classified into more than one biosynthetic pathway were grouped under their primary pathway classification and labeled as hybrids. (B) Relative abundance of terpenoid superclasses. (C) Relative abundance of amino acid and peptide superclasses. (D) Relative abundance of alkaloid superclasses. For each superclass, the intensities of metabolites within the respective superclass were grouped and summed. Metabolites classified into more than one superclass were grouped under their primary superclass classification and labeled as hybrids. NP = natural product

Within the terpenoid pathway (**Figure 1B**), sesquiterpenoids were the predominant superclass, underscoring their significance in the chemosystematics and bioactivity of *Ferula* species. Small peptides made up most of the metabolites from the amino acid and peptide metabolic pathway (**Figure 1C**), with the amino acids tyrosine, tryptophan, phenylalanine, and asparagine in high abundance (relative abundance > 1 %) (Table S1). The alkaloid biosynthetic pathway in *F. violacea* displayed extensive metabolite richness (**Figure 1D**), with tryptophan-, nicotinic acid-derived, and lysine-derived alkaloids being the most abundant superclasses. Highly abundant alkaloid metabolites (relative abundance > 1 %) included 3-indoleacrylic acid and 1-(6-methylpyridin-3-yl)ethanamine. The substantial presence of indole-based alkaloids is consistent with the observed expansion of tryptophan-derived metabolites, suggesting a strong reliance on aromatic amino acid metabolism for alkaloid biosynthesis.

**Figure 2** highlights the structural diversity and abundance of terpenoids in *F. violacea*. Several compounds were identified as highly conserved and abundant across *F. violacea* samples (**Table 5**). Among these, non-stereochemical parent compounds of nuciferol (CID: 78409369), fervanol vanillate (CID: 75226815), fetidone B (CID:73123203), a merosesquiterpenoid (CID: 74135949), and a group of structurally related daucane sesquiterpenoids exhibited the highest intensities.

**Figure 2.**
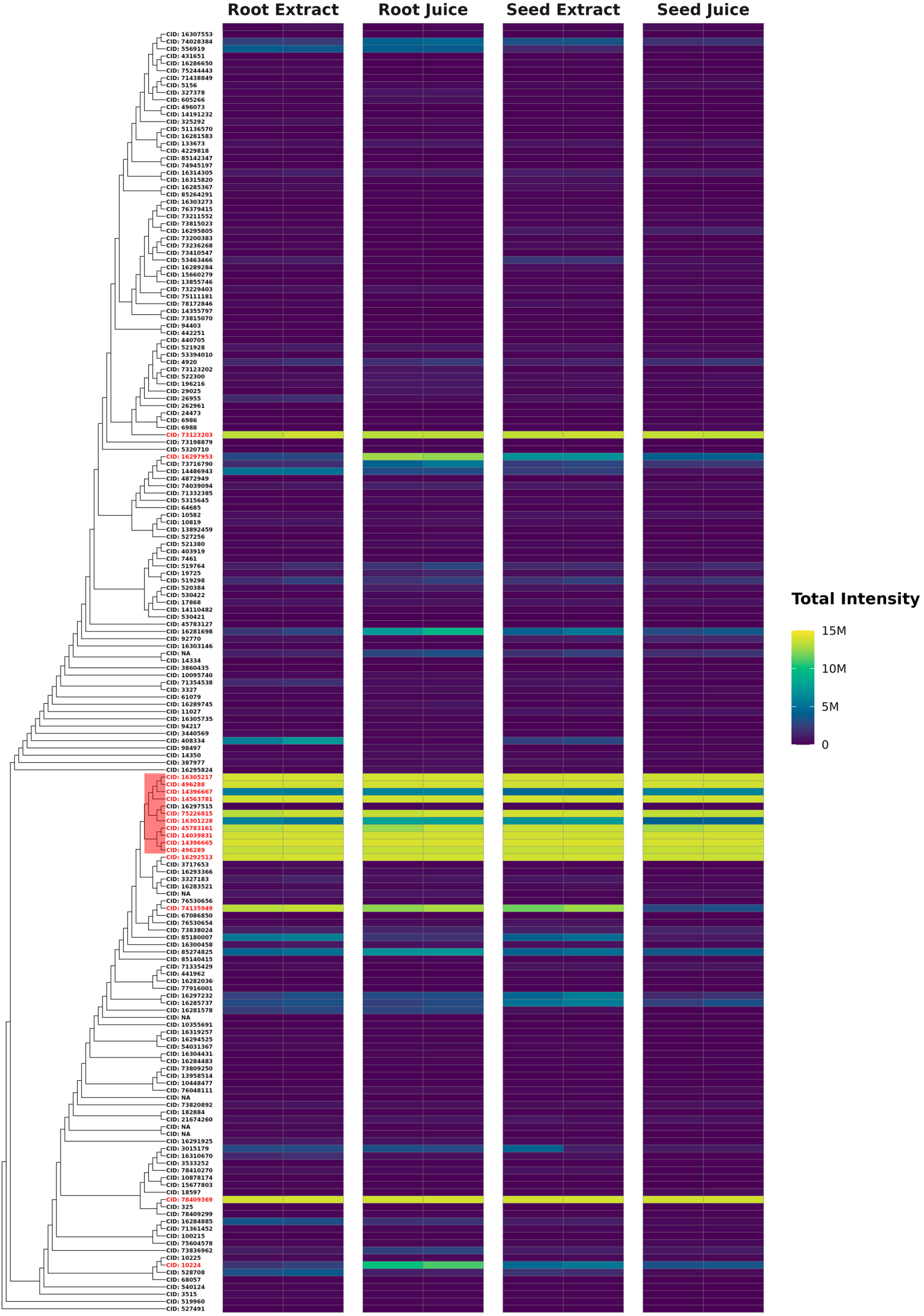
Abundance of Terpenoids Identified in *F. violacea*. Dendrogram and intensity heatmap of terpenoids in *F. violacea*, clustered by Tanimoto dissimilarity based on PubChem fingerprints. Leaf labels indicate the PubChem CID of each terpenoid (if available), with highly abundant terpenoids (relative abundance > 1 %) shown in red text. A group of structurally similar and highly abundant daucane sesquiterpenoids is highlighted in red. M = million

**Table 5.**
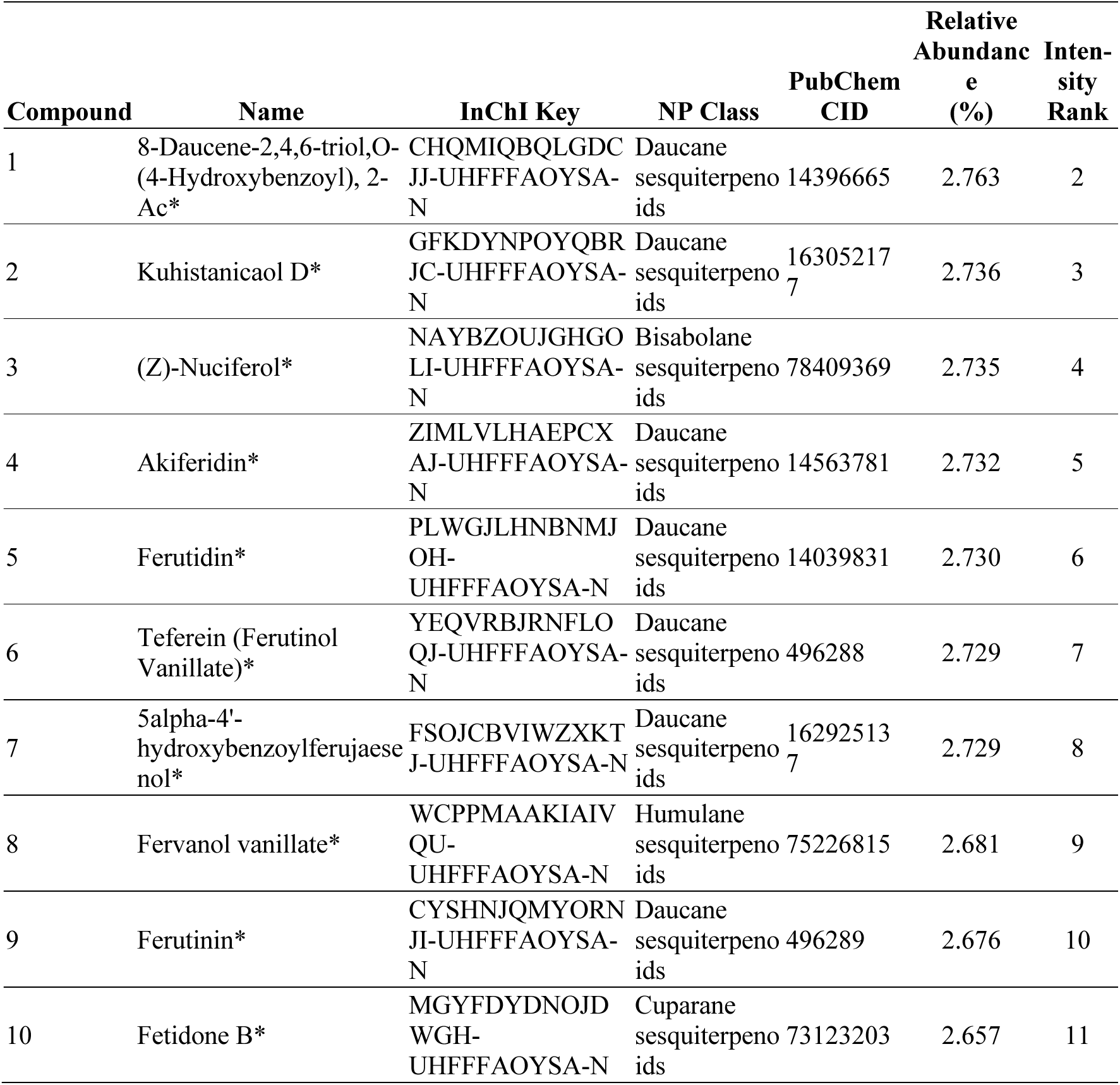
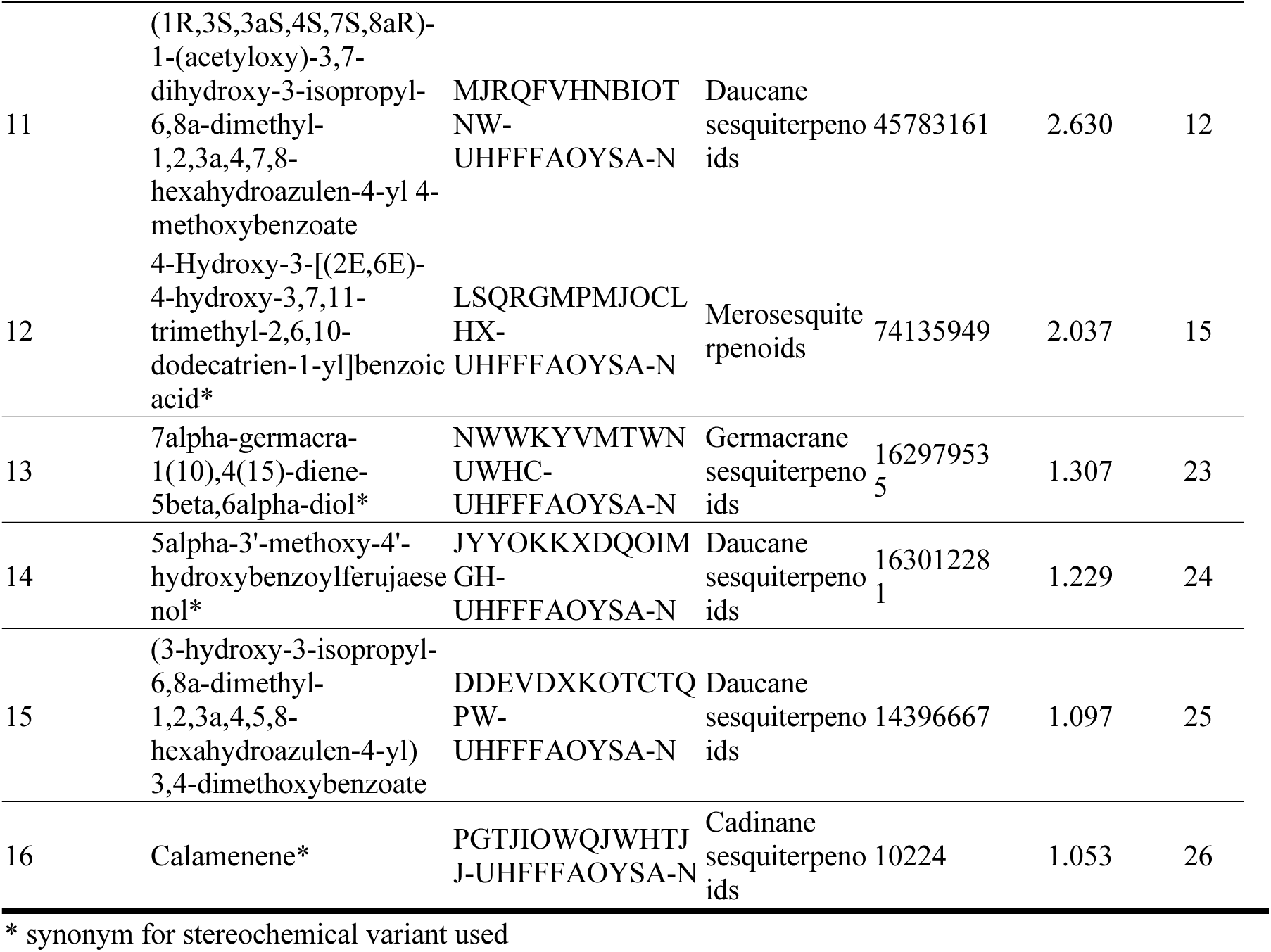
Top terpenoids putatively identified in *F. violacea*.

**Figure 3** provides a detailed structural analysis of the most abundant terpenoids identified in *F. violacea*, highlighting a conserved substructure and key functional modifications. The conserved (6-methyl-azulen-4-yl) cyclohexanecarboxylate substructure suggests a close biosynthetic relationship among these daucane sesquiterpenoids and may be a key determinant of their bioactivity. However, functional group modifications at positions 2, 3, 4, 12, 13, 14, and 15 introduce variability within the group. These modifications are likely to influence the biological activity, solubility, and interaction of these compounds with molecular targets, contributing to their diverse pharmacological properties. The high abundance and conservation of these compounds across samples suggest a strong biosynthetic preference for this class of compounds in *F. violacea* and potential pharmacological relevance.

**Figure 3.**
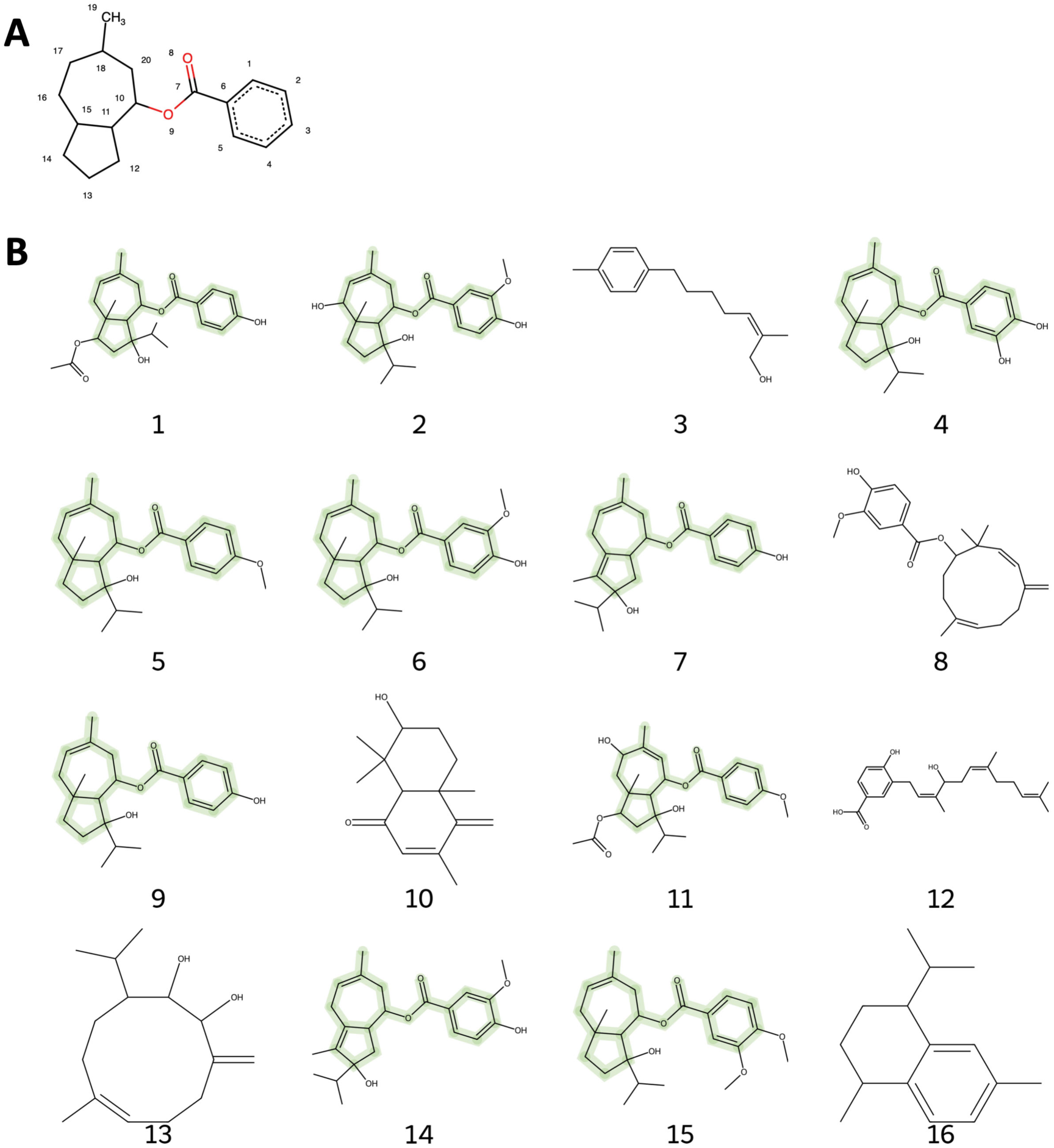
Structures of top terpenoids identified in *F. violacea* samples. (A) Common substructure of the highly abundant daucane sesquiterpenoid group. (B) Structures of top terpenoids (relative abundance > 1 %) generated from SMILES notation, with the common substructure highlighted in green. Structure identities and information are provided in Table 5.

### 2.3. Metabolic Differences Between *F. violacea* Roots and Seeds

The metabolite composition between *F. violacea* roots and seeds exhibited significant differences, as demonstrated by Principal Component Analysis (PCA) (**Figure 4A**) and PERMANOVA (variable: organ, R2: 35 %, p-value: 0.005). A total of 532 metabolites were identified in root samples, with 21 being exclusive to roots, while 519 were detected in seed samples, of which only 8 were unique to seeds (**Figure 4B**). The classification of organ-specific metabolites indicates that metabolites unique to roots and seeds are primarily associated with the shikimate and phenylpropanoid, terpenoid, and alkaloid biosynthetic pathways (**Figure 5**). The majority of these compounds were detected at relatively low abundance, with signal intensities below 1 million. In addition, metabolite differences between organs were dependent on the sample processing method as shown by PCA (**Figure 4A**) and confirmed by PERMANOVA (variable: organ+processing method, R2: 30 %, p-value: 0.013). Notably, root-specific metabolites included a distinct group of flavonoids and phenolic acids from the shikimate and phenylpropanoid pathway, which were exclusively detected in root juice samples (**Figure 5**).

**Figure 4.**
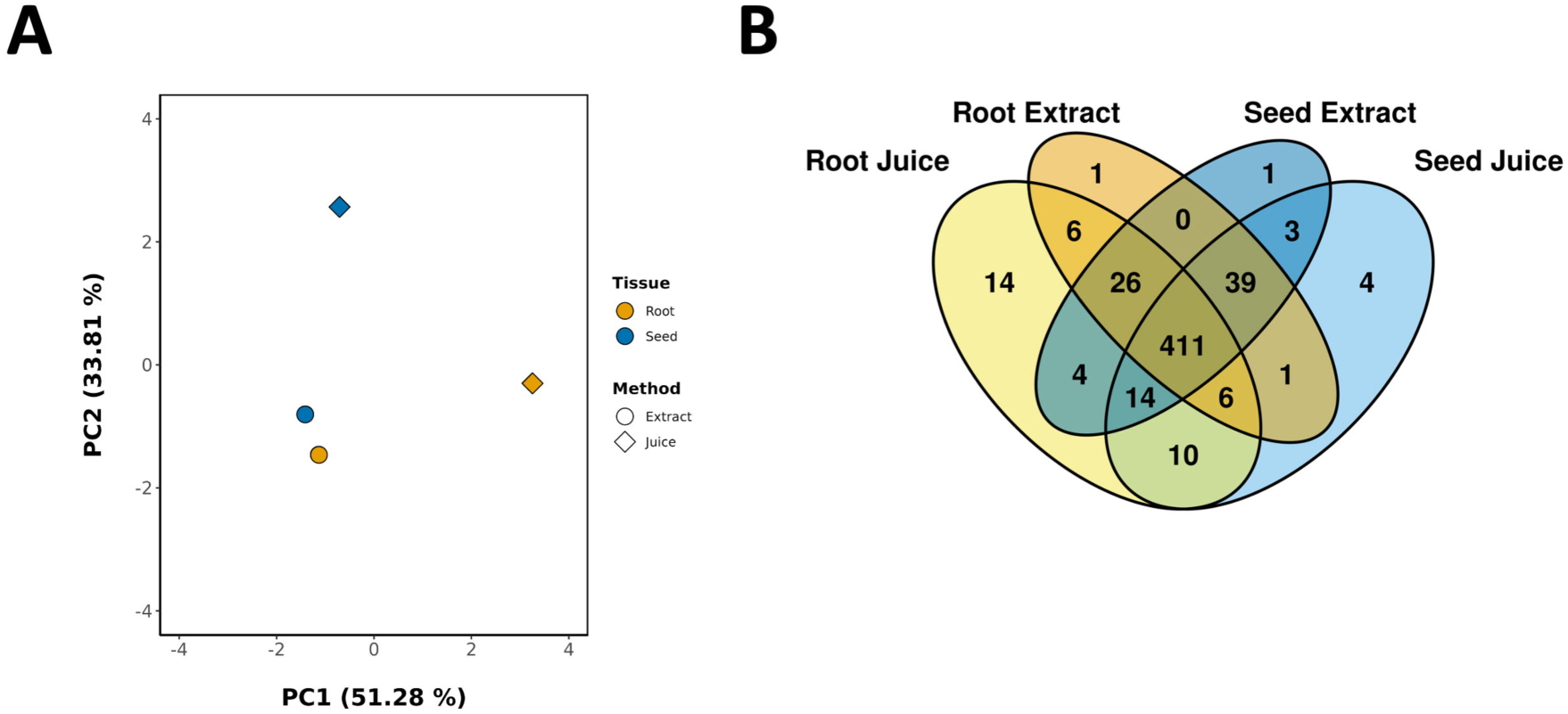
Global Analysis of Metabolites from *F. violacea* Samples. (A) Principal Component Analysis (PCA) of metabolite data. (B) Venn diagram showing the distribution of metabolites between samples.

**Figure 5.**
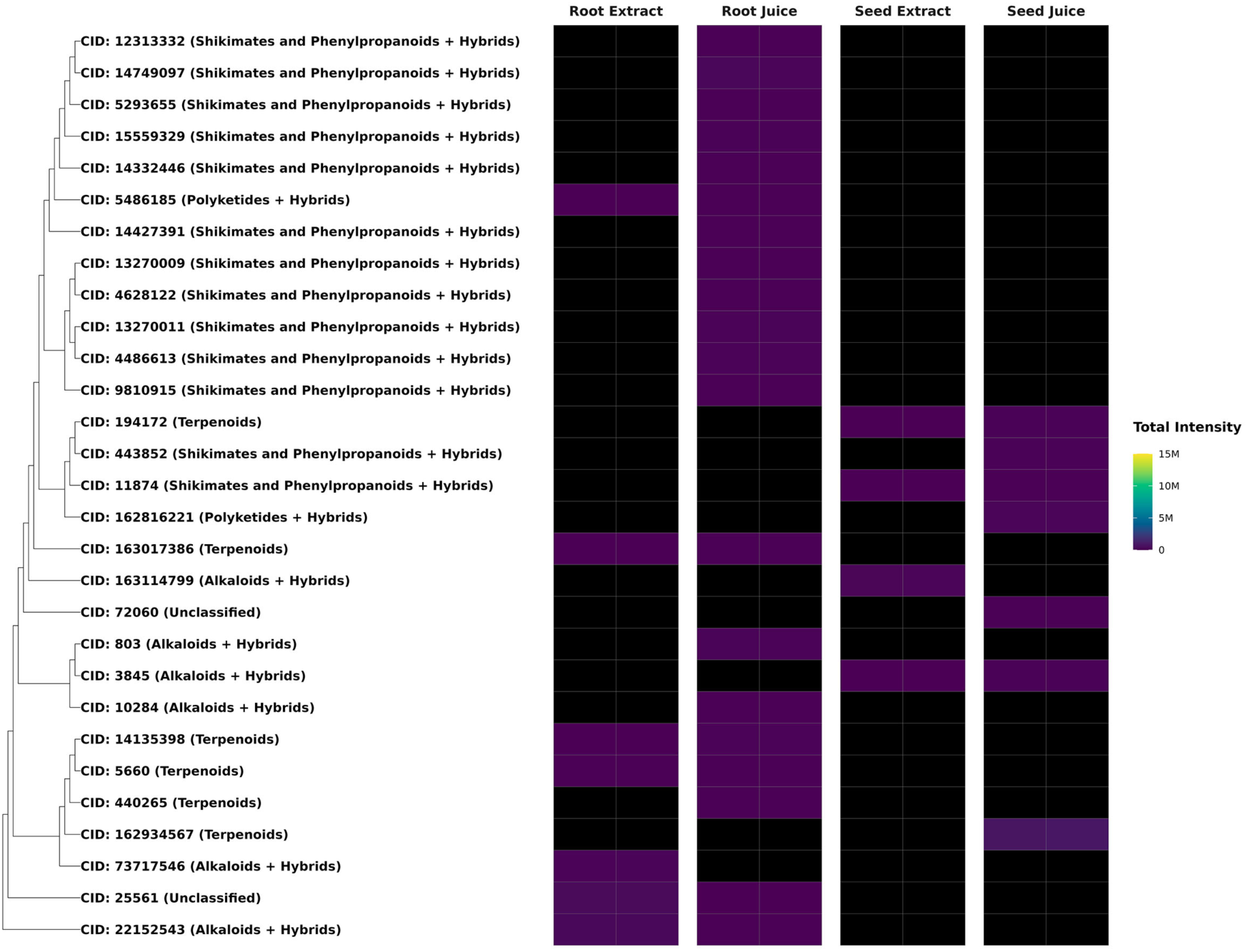
Organ-Specific Metabolites in *F. violacea* Roots and Seeds. Dendrogram and intensity heatmap of organ-specific metabolites clustered by Tanimoto dissimilarity based on PubChem fingerprints. Leaf labels indicate the PubChem CID of each metabolite (if available), with natural product biosynthetic pathway classification shown in parentheses. Black heatmap tiles indicate metabolites that were not detected in the respective sample (intensity < 10,000). M = million

Differential abundance analysis identified a total of 60 root-enriched and 83 seed-enriched metabolites (log2-fold change > 1, FDR < 0.05) (**Table S2**). Most metabolites with significant differences in abundance between organs (FDR < 0.05) exhibited log2-fold changes below 5, though a few metabolites displayed log2-fold changes above 8 (**Figure 6**), which included the seed-enriched terpenoid farnesyl-4-hydroxybenzoesaure (CID: 54248366). Other seed-enriched metabolites with intensities exceeding 1 million were predominantly alkaloids and amino acids (**Figure 7**). This suggests that seeds prioritize alkaloid biosynthesis, likely for chemical defense or germination-associated metabolic functions. In contrast, root-enriched metabolites with intensities exceeding 1 million were primarily terpenoids (**Figure 7**). These findings suggest organ-specific metabolic adaptation, potentially associated with root defense mechanisms or specialized biosynthetic functions.

**Figure 6.**
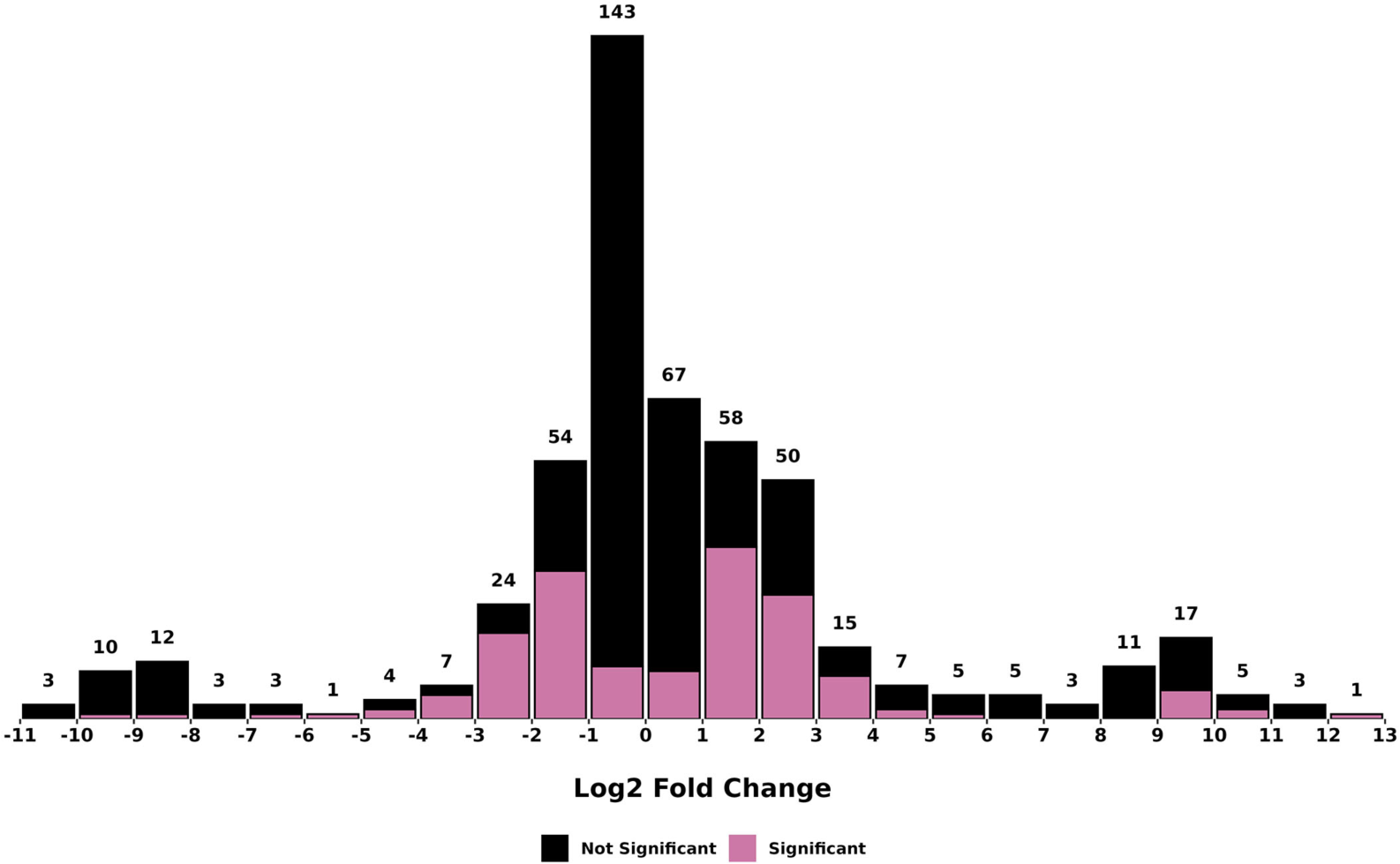
Histogram of log2-fold changes (seed/root) in metabolite abundance between *F. violacea* roots and seeds. The text on top of each bar indicates the total number of metabolites found in each bin. Metabolites with significant differences in abundance (FDR < 0.05) between organs are colored pink, while those with no significant differences are colored black.

**Figure 7.**
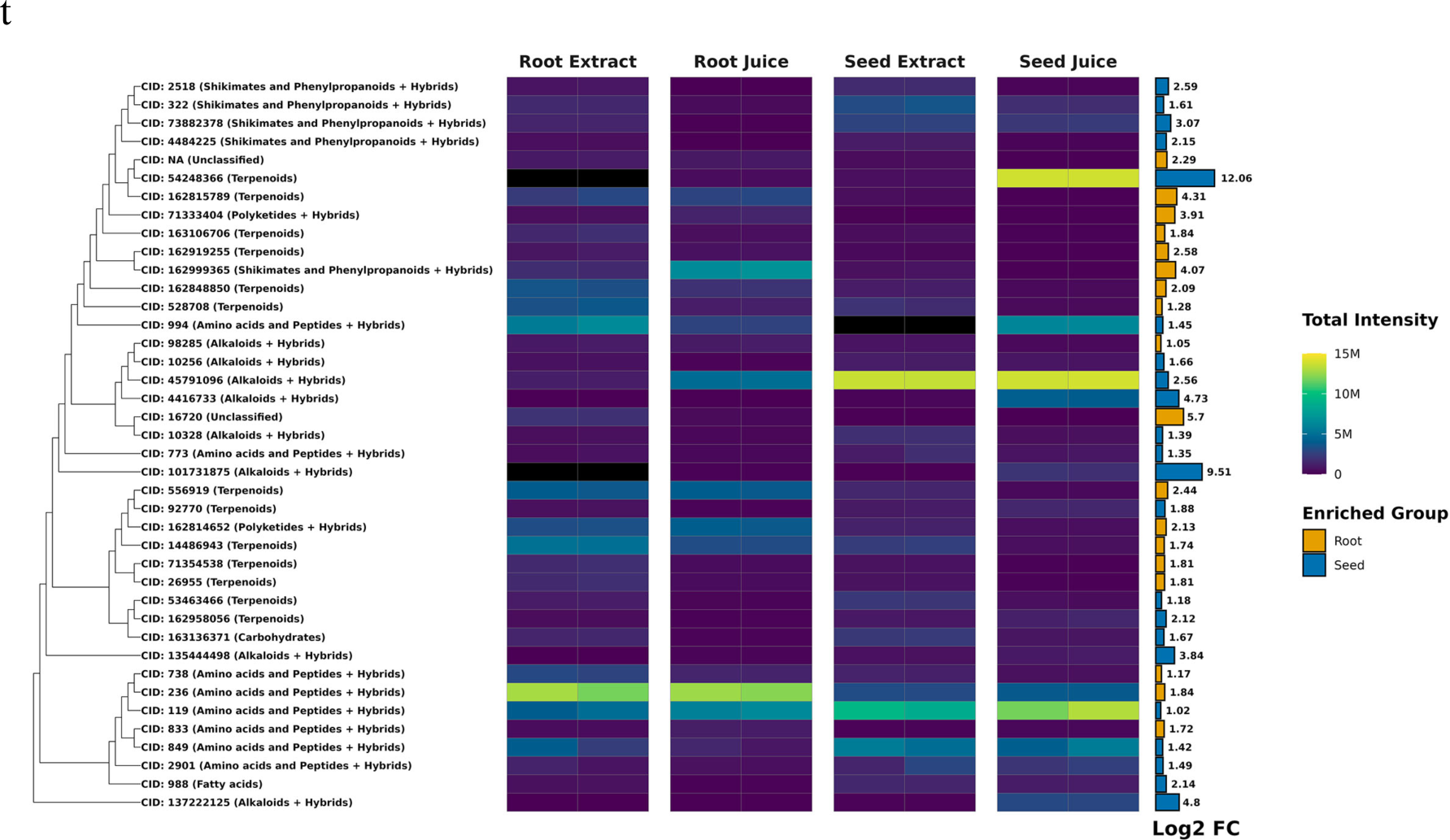
Differentially Abundant Metabolites Between Roots and Seeds of *F. violacea*. Dendrogram and intensity heatmap of differentially abundant metabolites (log2-fold change > 1 and FDR < 0.05) between organs with intensities greater than 1 million. The dendrogram was constructed using Tanimoto dissimilarity based on PubChem fingerprints. Leaf labels indicate the PubChem CID of each metabolite (if available), with natural product biosynthetic pathway classification shown in parentheses. Black heatmap tiles indicate metabolites that were not detected in the respective sample (intensity < 10,000). A bar graph of log2-fold change is displayed to the right of the heatmap, with colors representing the group in which each compound is enriched. M = million

### 2.4. Impact of Sample Processing Method on *F. violacea* Metabolite Composition

Metabolite composition was compared between juice and extract preparations to evaluate how sample processing influenced metabolite recovery. PCA showed clear separation between juices and extracts, with PERMANOVA also indicating that the sample processing method significantly impacted metabolite composition (variable: processing method, R2: 35 %, p-value: 0.009) (**Figure 4A**). In total, 512 metabolites were detected in ethanol extracts, including 2 unique to extracts, whereas 538 total metabolites were detected in juices, with 28 metabolites specific to juices (**Figure 4B**). This indicates that juicing captured almost all the metabolite diversity in *F. violacea*. Most juice-specific metabolites were associated with the shikimate and phenylpropanoid pathway and terpenoids. These method-specific compounds generally exhibited low abundance, with signal intensities below 3.0 million (**Figure 8**). A recurring group of structurally related shikimate and phenylpropanoid derivatives from the flavonoid and phenolic acid superclasses was detected exclusively in root juice samples, suggesting preferential solubilization of these metabolites in aqueous extracts.

**Figure 8.**
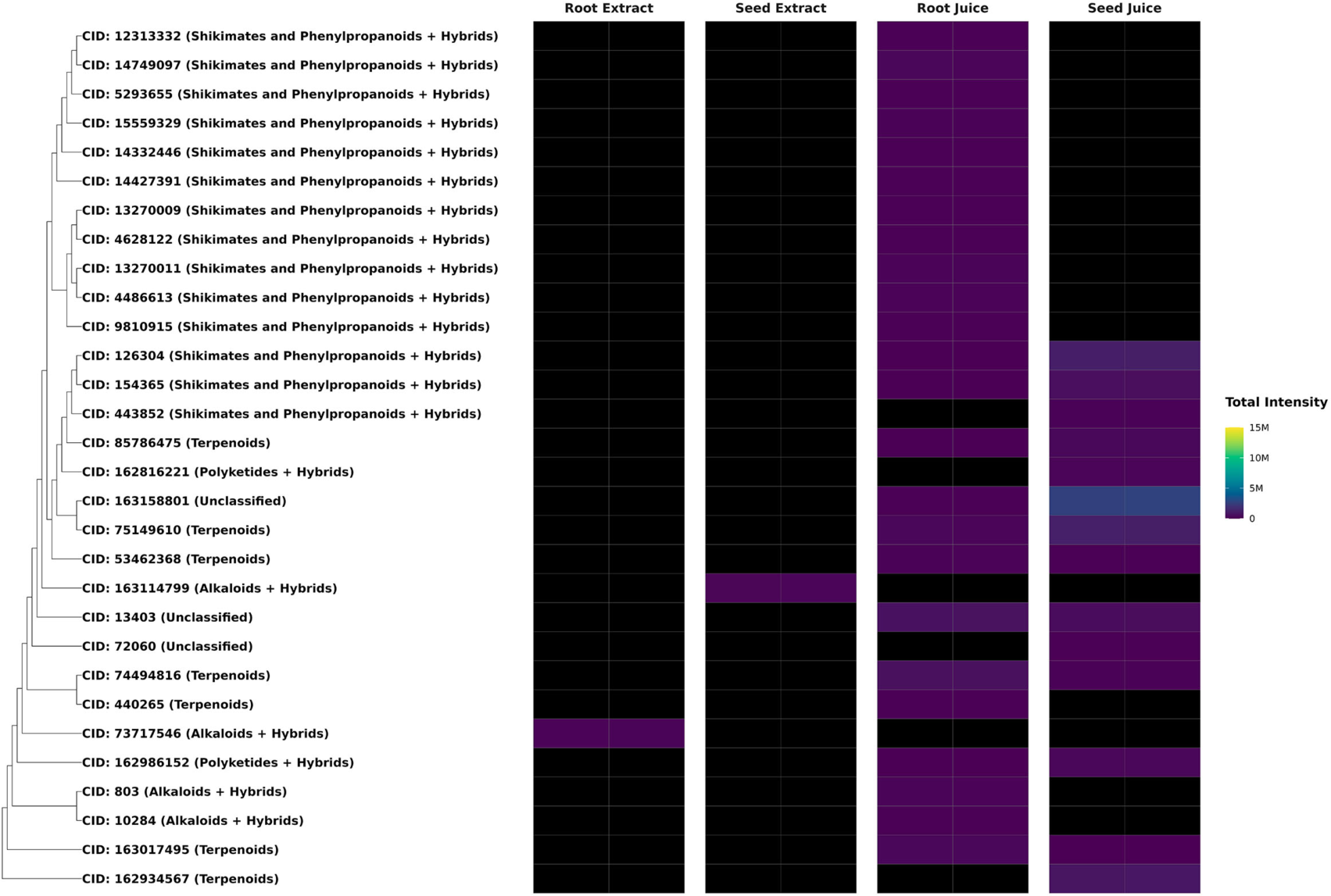
Metabolites Specific to *F. violacea* Extracts and Juice. Dendrogram and intensity heatmap of processing method-specific metabolites clustered by Tanimoto dissimilarity based on PubChem fingerprints. Leaf labels indicate the PubChem CID of each metabolite (if available), with natural product biosynthetic pathway classification shown in parentheses. Black heatmap tiles indicate metabolites that were not detected in the respective sample. (intensity < 10,000) M = million

Differential abundance analysis identified 65 metabolites enriched in extracts and 32 enriched in juices (**Table S3**). Many differentially abundant metabolites between processing methods exhibited a log2-fold change of less than 5 (**Figure 9**). Nearly all highly abundant differentially enriched metabolites (intensities > 1 million) were found in extracts (**Figure 10**). Notably, extracts yielded a greater number of enriched terpenoids with intensities exceeding 1 million, highlighting the distinct chemical profiles associated with different extraction methods. In juices, the terpenoid farnesyl-4-hydroxybenzoesaure (CID: 54248366) was highly enriched and abundant specifically in seed juices. These metabolic distinctions underscore the impact of sample processing methods on metabolite composition and the specialized biosynthetic capacities of each organ, likely reflecting their distinct physiological and ecological functions.

**Figure 9.**
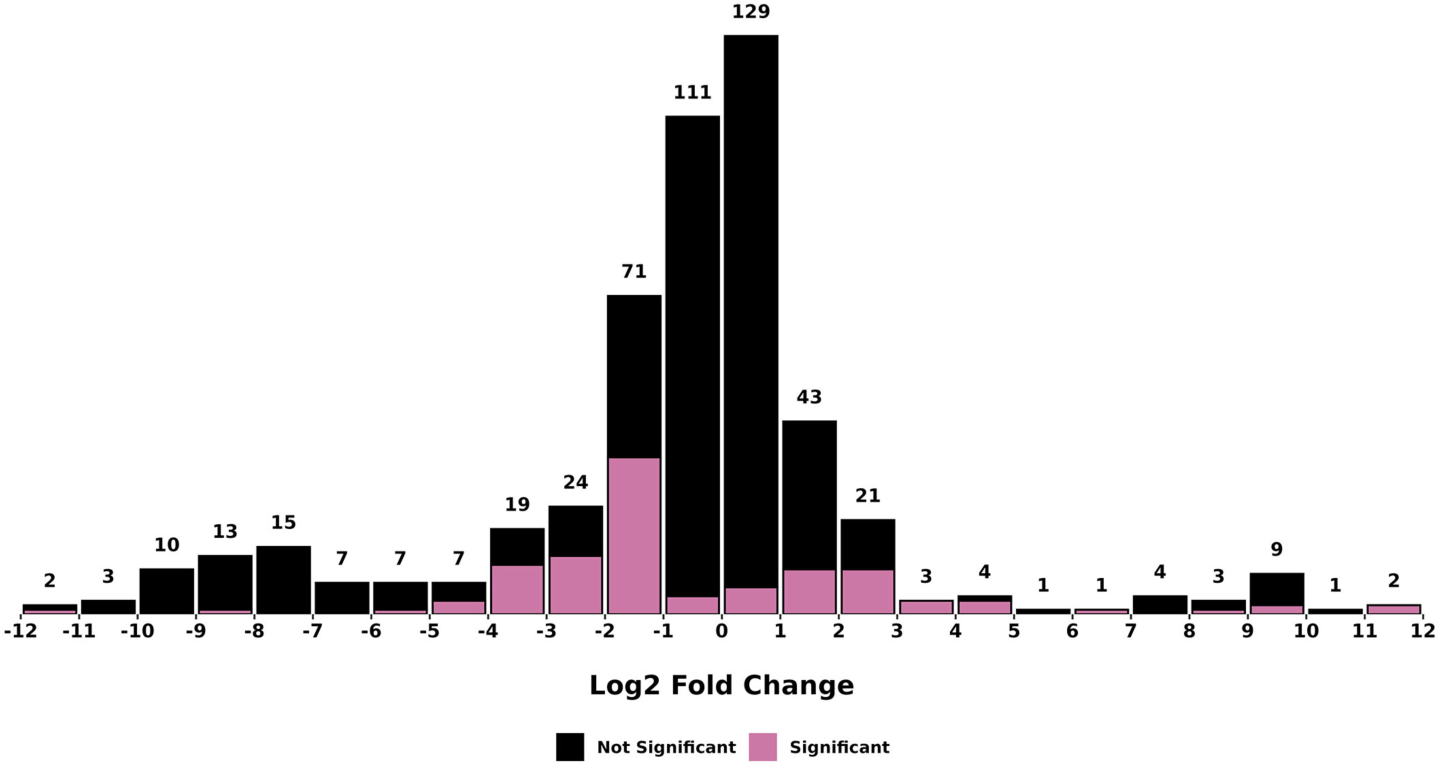
Histogram of log2-fold changes (juice/extract) in metabolite abundance between *F. violacea* processing methods. The text on top of each bar indicates the total number of metabolites found in each bin. Metabolites with significant differences in abundance (FDR < 0.05) between processing methods are colored pink, while those with no significant differences are colored black.

**Figure 10.**
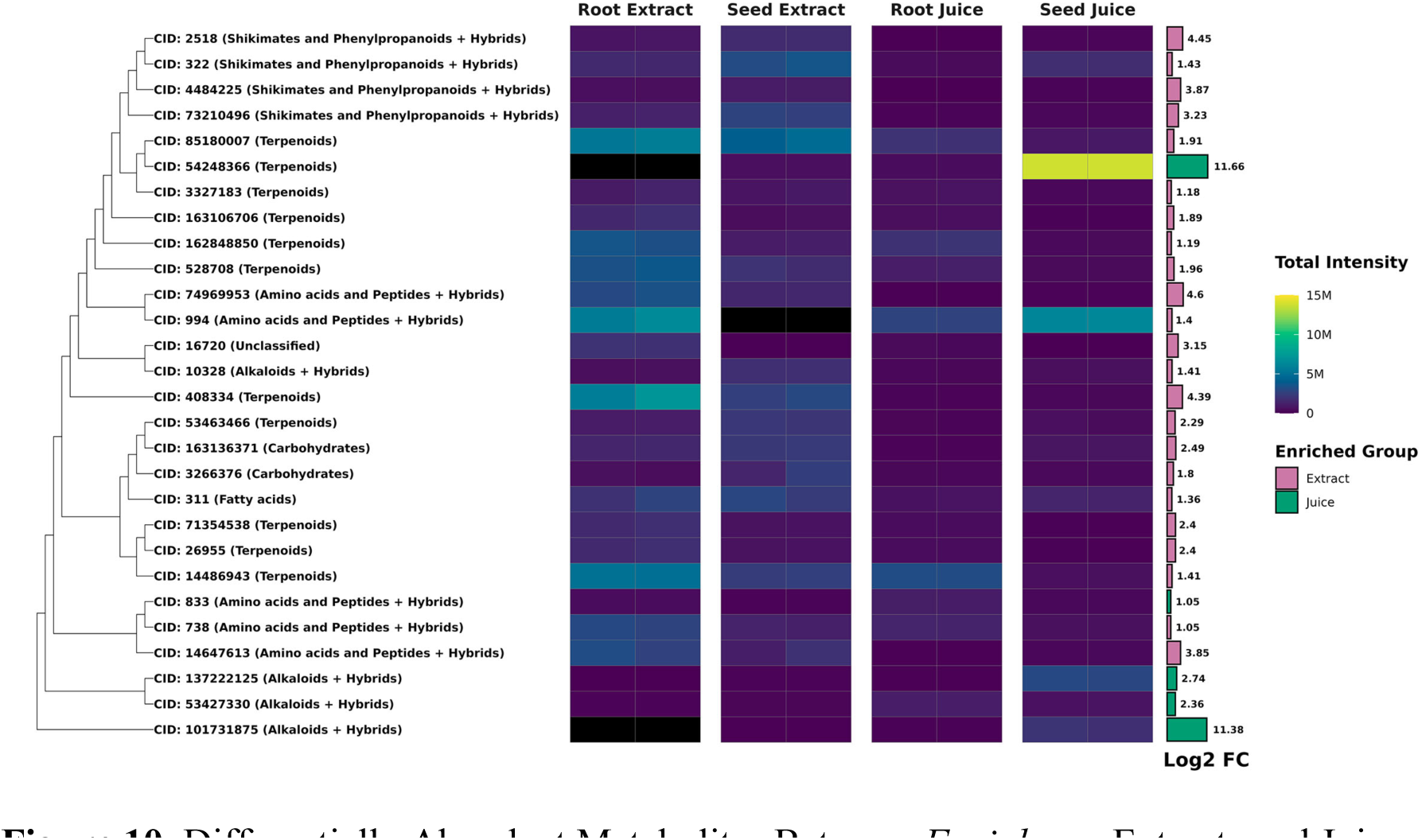
Differentially Abundant Metabolites Between *F. violacea* Extracts and Juices. Dendrogram and intensity heatmap of differentially abundant metabolites (log2-fold change > 1 and FDR < 0.05) between sample processing methods with intensities greater than 1 million. The dendrogram was constructed using Tanimoto dissimilarity based on PubChem fingerprints. Leaf labels indicate the PubChem CID of each metabolite (if available), with natural product biosynthetic pathway classification shown in parentheses. Black heatmap tiles indicate metabolites that were not detected in the respective sample (intensity < 10,000). A bar graph of log2-fold change is displayed to the right of the heatmap, with colors representing the group in which each compound is enriched. M = million

## 3. Discussion

### 3.1. Bioactive metabolites in *Ferula* species

The medicinal properties of *Ferula* species are largely attributed to their oleo-gum resins, which are harvested by cutting the stem sprout in spring and periodically collecting the exudate. Sulfur-containing metabolites, such as (E)-1-propenyl-sec-butyl disulfide and (Z)-1-propenyl-sec-butyl disulfide, are responsible for the pungent odor of these species and possess antimicrobial, antifungal, and carminative properties, making them valuable in treating gastrointestinal and respiratory disorders. Recent research also suggests their potential to modulate oxidative stress pathways (Elarabany, Hamad et al. 2023, Jiang, Peng et al. 2024). Additional volatile compounds, including monoterpenes and sesquiterpenes such as α-pinene, β-eudesmol, limonene, and myrcene, are found in the essential oils of *Ferula* (Attarian, Kavoosi et al. 2024, Ghaforzoda, Sharopov et al. 2025). These oils also contribute to the fragrant odors of many *Ferula* species and exhibit significant antimicrobial, antifungal, and insecticidal activities, highlighting their potential in developing sustainable pest management solutions. The widespread occurrence of terpenoids, including diterpenes and sesquiterpenes, and their associated antimicrobial, antifungal, and antiparasitic effects, further underscores the therapeutic potential of this genus (Abaza 2024).

Coumarins, particularly sesquiterpene coumarins including umbelliprenin and badrakemin, are hallmark constituents of *Ferula* species (Khosnutdinova, Gemejiyeva et al. 2023). Coumarins are one of the most common organic molecules and are used in medicine for their pharmacological effects, including anti-inflammatory, anticoagulant, antihypertensive, anticonvulsant, antioxidant, antimicrobial, and neuroprotective (Flores-Morales, Villasana-Ruíz et al. 2023). Isolated from species such as *F. gummosa* Boiss. and *F. assa-foetida*, these compounds have demonstrated cytotoxic activity against various cancer cell lines, signifying their potential in anticancer drug development (De Luca, Salim et al. 2012, Elarabany, Hamad et al. 2023). Furthermore, sesquiterpenes and their derivatives, including sesquiterpene phenylpropanoids and sesquiterpene chromones, have garnered scientific interest due to their diverse chemical structures and promising biological properties, including antioxidative, anti-inflammatory, and antibacterial activities (Wang, Zheng et al. 2023).

*Ferula* species are rich in phenolic compounds. Flavonoids and phenolic acids contribute significantly to their antioxidant capabilities and a broad spectrum of pharmacological activities. These phenolic compounds are known for their health-promoting effects, including anticancer, anti-inflammatory, antioxidant, antimicrobial, and anti-aging properties (Nouioura, El Fadili et al. 2024). Studies using HPLC–DAD have identified bioactive substances in the non-volatile fraction of *F. communis*, likely responsible for these effects. The presence of flavonoids like quercetin and kaempferol in several *Ferula* species supports their traditional use in managing inflammatory and cardiovascular conditions (Sharopov and Setzer 2018). The notable bioactivity of the *Ferula* species underscores the importance of comprehensive phytochemical investigations.

### 3.2 Expansion of *Ferula* metabolite diversity

The phytochemical content and bioactivity of *Ferula* species can vary significantly. For example, research on *F. assa-foetida* has revealed differences in phenolic content, antioxidant activity, and essential oil profiles among populations growing at different elevations, with higher antioxidant activity and phenolic content observed in populations at higher altitudes (Nouioura, El Fadili et al. 2024). Continued elucidation of these compounds through advanced analytical techniques holds promise for the discovery and development of novel pharmaceuticals with the potential to improve human health outcomes. In this study, the untargeted metabolomic analysis of *F. violacea* significantly expanded the known chemical diversity of the *Ferula* genus. This study identified 540 distinct metabolites in *F. violacea*. Of these, 536 are reported for the first time in this species, and 419 have not been previously reported in the genus *Ferula* (Table S1, **Figure 1**). This large number of newly reported metabolites may be attributed to the limited number of LC-MS/MS untargeted metabolomics studies on *Ferula*, resulting in gaps in these databases. This bias is demonstrated by the finding that 416 out of the 419 newly reported metabolites can be found in other species (Table S1). Additional bias in natural product databases can arise from their specific focus and lack of regular updates, as evident in databases such as NPASS, which is limited to experimentally determined bioactive natural products and was last updated in 2023. With respect to previous LC-MS/MS untargeted metabolomics studies of *Ferula*, differences in sampling, extraction, and analytical procedures may explain the discrepancies in detected metabolites, with in-silico MS/MS fragmentation analysis (SIRIUS) in this study likely enhancing metabolite identification.

Our results significantly increase the number of reported natural products across multiple biosynthetic pathways. The most significant chemical expansion was observed in the terpenoid, shikimate and phenylpropanoid, and alkaloid pathways. Among the 143 newly identified terpenoids (Table 1), sesquiterpenoids and monoterpenoids were the most abundant subclasses, with a particular dominance of daucane-type sesquiterpenoids (Table 2). Sesquiterpenes are a subclass of terpenes, which are natural hydrocarbons synthesized by plants, including the *Ferula* species, through the mevalonic acid pathway. Sesquiterpenes and their derivatives have antibacterial, antifungal, and antiviral activities, which are characteristically related to plant defense mechanisms (Wang et al., 2023). These compounds are known for their pharmacological properties, including antimicrobial and cytotoxic activities, suggesting that *F. violacea* may serve as a valuable source of bioactive terpenoids.

**Figure 3** illustrates the variety of daucane sesquiterpenoids identified in *F. violacea*, demonstrating that these metabolites constitute the most abundant group of secondary metabolites in this species (Table 5), approaching 25% in relative abundance. Similarly, other *Ferula* species have been identified as major sources of naturally occurring daucane-type sesquiterpenoids. Ferutinin, a daucane-type sesquiterpene ester, was first described in the early 1970s from *F. tenuisecta* Eug. Kor. (Saidkhodzhaev and Nikonov 1973). It is a characteristic compound for the genus *Ferula*, identified in multiple species (Wang, Zheng et al. 2023). As demonstrated in this study, ferutinin is also a prominent metabolite in *F. violacea*, reaching up to 2.7% of the relative abundance of identified phytochemicals. Baykan and colleagues (Baykan, Aydogan et al. 2020) evaluated Ferutinin content in the roots of 12 *Ferula* species growing in Turkey. They found that the ferutinin was present in high concentrations in the n-hexane extracts compared to the chloroform extracts and for six species (*F. communis, F. huber-morathii, F. orientalis, F. rigidula, F. szowitsiana, F. tingitana*). Ferutinin was found only in the n-hexane extract and was absent in *F. lycia* and *F. drudeana*, demonstrating its dependence on species and extraction method. The highest concentration of the compound was found in *F. tenuissima* and *F. halophila* n-hexane extracts (167,380 µg/g and 157,160 µg/g, respectively) (Baykan, Aydogan et al. 2020). Daucane-type sesquiterpenes play a major role in the bioactivity of *Ferula* and are responsible for the broad use of this plant’s health-related applications. For example, ferutinin has been reported as a phytoestrogen acting as an agonist for ER-α and an agonist/antagonist for ER-β [14,(Ikeda, Arao et al. 2002) and to have anticancer activity (Aydoğan, Baykan et al. 2019).

The shikimate and phenylpropanoid pathway also experienced a substantial increase in novel chemical diversity with *F. violacea*, with 83 previously unreported metabolites. The shikimate pathway is the metabolic process responsible for the biosynthesis of the aromatic amino acids (Shende, Bauman et al. 2024) that are common in *F. violacea*. This class markedly included notable expansions in coumarins, flavonoids, and lignans, which are widely recognized for their antioxidant and anti-inflammatory activities (Hassanein, Althagafy et al. 2024, Younes and Mustafa 2024, Zheng, Zhang et al. 2025). It is important to emphasize the documented anticoagulant activity of coumarins (Younes and Mustafa 2024), a broad spectrum of cardiovascular (Ciumărnean, Milaciu et al. 2020, Das, Keerthi et al. 2025) and neuroprotective (Bellavite 2023, Mishra 2024) properties of flavonoids along with the estrogen-like activity of lignans (Li, Ma et al. 2023, Bagheri and Esmailidehaj 2024). The presence of small peptides and diazotetronic acid derivatives suggests a possible interaction between phenylpropanoid metabolism and non-ribosomal peptide biosynthesis, further broadening the chemical landscape of *F. violacea*.

A particularly noteworthy expansion was observed in alkaloids, where 56 metabolites were previously unreported in *Ferula*. Alkaloids, naturally occurring compounds in a diverse range of plant species, hold vast potential for biological, medicinal, and pharmacological applications (Schläger and Dräger 2016, Heinrich, Mah et al. 2021). The identification of tetramate and peptide alkaloids suggests the presence of non-ribosomal peptide synthetase (NRPS) pathways, an uncommon feature in *Ferula* species that may have biotechnological applications. *Ferula* plants have inhibitory effects on various viruses, making them an attractive alternative to conventional antiviral agents. Therefore, these plants are a natural source of valuable compounds that can help fight infectious diseases (Mohammadi, Forouzanfar et al. 2025). It is also important to mention the synergistic potential of alkaloids when combined with other phytochemicals, offering new insights into more potent, multi-compound therapeutic formulations (Letchuman, Madhuranga et al. 2024, Das and Ruhal 2025). This may contribute to antiviral, antibacterial, and antifungal properties recently reported for *F. violacea* (Satorov, Mavlonazarova et al. 2024, Satorov, Mavlonazarova et al. 2024).

The phytochemical differences between plant roots and seeds are substantial, reflecting their specialized functions within the plant (Arumugam, Elanchezhian et al. 2022). Significant differences were identified between the metabolite profiles of *F. violacea* roots and seeds. The root metabolome was enriched in shikimate and phenylpropanoid derivatives, which may contribute to lignin biosynthesis and root defense mechanisms. In contrast, seeds contained a higher abundance of alkaloids and amino acids, potentially reflecting their role in chemical defense and germination-associated metabolism. The organ-specific distribution of these metabolites indicates that secondary metabolism in *F. violacea* is highly specialized and functionally adaptive.

The method of extraction significantly affects the availability of bioactive compounds (Baimakhanova, Sadanov et al. 2025). It has been shown that ethanol extracts are often particularly rich in total flavonoids, whereas aqueous extracts contain higher levels of phenolic compounds (Sahoo, Raghava Varrier et al. 2024). The method of sample processing significantly influenced the chemical composition of *F. violacea* extracts. Juice samples were notably enriched in shikimate and phenylpropanoid metabolites, particularly in root tissues, suggesting that these metabolites are less soluble in ethanol extractions. In contrast, ethanol extracts contained a greater number of terpenoids and amino acids, highlighting the role of solvent polarity in the selective recovery of bioactive compounds (Yadav, Patel et al. 2025). Juice samples also captured nearly the full metabolite diversity of *F. violacea*, containing 538 out of 540 detected metabolites, compared to 512 found in ethanol extracts.

### 3.3 Broader implications

This study substantially expands the known metabolite repertoire of *F. violacea* and the genus *Ferula*, adding 419 metabolites to its phytochemical inventory. The prevalence of daucane sesquiterpenoids consolidates their role as chemotaxonomic markers, while the unexpectedly broad alkaloid diversity indicates potentially novel biosynthetic pathways and merits genetic and biochemical investigation. Organ- and method-specific patterns underline the importance of sampling strategy in metabolomics studies.

While our analysis was restricted to UHPLC-QTOF-MS and did not include orthogonal confirmation by GC-MS or NMR, the integrated use of spectral libraries, cheminformatics, and database cross-referencing provides high-confidence structural annotations. Future targeted studies with authentic standards will be necessary to validate structural isomers and quantify pharmacologically relevant constituents.

## 4. Materials and methods

### 4.1 Plants

*Ferula violacea* Korovin plants were collected on July 23, 2022, from the Maykhura region in Tajikistan, located approximately 67 km north of Dushanbe in the Region of Republican Subordination (Latitude: 38.48319, Longitude: 68.490523, Elevation: 1180 MASL). Plant specimens were identified utilizing herbarium sheets from Moscow University (Seregin 2023), and subsequently verified by staff at the Botanical Garden of the Academy of Sciences of Tajikistan. The species name has been validated and is accepted by the World Flora Online (WFO 2024).

### 4.2 Sample preparation

The roots were thoroughly washed, dried, and finely chopped using a scalpel. The prepared plant material was submerged in 70% ethanol at a ratio of 100 g of root to 100 mL of ethanol. Maceration was performed at room temperature for 24 hours. Following maceration, the mixture was processed using a juicer (Model SPV-2, manufactured in Kharkiv, 1984) to enhance extraction. The resulting extract was collected in Petri dishes and dried in a vacuum oven (Drier Box DHG-9053A) at 40–45°C for 24 hours. Dried extracts were stored in airtight vials for further analysis. In a separate process, roots were directly juiced, and the resultant liquid was collected in Petri dishes, dried in a vacuum oven at 40–45°C for 24 hours, and subsequently stored in airtight vials. Additionally, roots were processed using a juicer. The juice was collected, dried in a vacuum oven at 40–45°C for 24 hours, and stored in airtight vials for subsequent use.

For seed processing, seeds were bisected and placed in glass or porcelain vessels. One hundred grams of seeds were submerged in 100 mL of 70% ethanol and macerated at room temperature for 24 hours. The mixture was filtered through a 1 mm sieve to remove solids, and the filtrate was dried in a vacuum oven at 40–45°C for 24 hours. Dried extracts were stored in airtight vials. Additionally, seeds were processed using a juicer. The juice was collected, dried in a vacuum oven at 40–45°C for 24 hours, and stored in airtight vials for subsequent use.

### 4.3 Phytochemical analysis

Samples were diluted in a 50:50 mixture of methanol and water, followed by sonication using a Bioruptor sonication system. Analytical measurements were performed using a Bruker Daltonics maXis-II UHR-ESI-QqTOF mass spectrometer coupled with a Thermo Scientific Ultimate-3000 UHPLC system. Up to 10 μL of sample was injected onto an Agilent Acclaim 120 C18 column (2.1 mm × 150 mm, 2.2 μm) maintained at 30 °C with a flow rate of 150 μL/min.

The gradient elution program consisted of 2% solvent B (acetonitrile with 0.15% formic acid) and 98% solvent A (water with 0.15% formic acid) for the first 2 minutes, followed by a gradient increase to 40% solvent B over 20 minutes, 98% solvent B over the next 10 minutes, and held at 98% solvent B for an additional 10 minutes. Mass spectrometry data were acquired over an m/z range of 50–1300 using positive ion mode electrospray ionization. Raw data were processed using MetaboScape 2024b software (Bruker Daltonics) alongside multiple metabolomics databases: the Bruker MetaboBASE Personal Library 3.0 (), MassBank of North America (MoNA) with LipidBlast2022 (), and COCONUT () (Chandrasekhar, Rajan et al. 2025). In-silico fragmentation analysis was conducted using integrated MetaboScape tools and SIRIUS 6.2.2 software with CSI:FingerID scoring (Dührkop, Fleischauer et al. 2019, Hoffmann, Nothias et al. 2022).

### 4.4 Processing of mass feature table

Data harmonization was performed on the MetaboScape mass feature table. For features lacking InChI notations but containing MetaboScape compound names, we retrieved corresponding InChI key identifiers from PubChem (NIH, 2024) using PubChemPy (version 1.0.4)(Swain 2014). Conversely, for features with InChI notations but missing compound names, we conducted reverse searches to obtain standardized chemical nomenclature.

Unknown molecular features were identified according to these criteria:1) Library matches: MoNA or MetaboBASE Personal Library matches with MS/MS scores >600; 2) Computational predictions: COCONUT or HMDB matches with in-silico fragmentation scores >800, and/or SIRIUS 6.2.2 analysis yielding top-confidence matches; 3. Spectral quality: mSigma values <30 and mass accuracies within ±2.5 ppm.(Dührkop, Fleischauer et al. 2019). Missing values were imputed by substituting 0 for absent features across all replicates or by replacing them with the minimum intensity minus 1 for features missing in some replicates. Noise removal excluded features present only in blank samples, those with higher intensities in blank samples, or those with maximum intensities below 20,000. The median intensities of blank samples were subtracted from the intensities of experimental samples to finalize the mass feature table.

### 4.5 Retrieval and Annotation of Natural Products from *Ferula*

Natural products reported in the *Ferula* genus were retrieved from multiple sources, including the Reaxys chemical database (Reaxys 2024)(accessed May 28, 2025), a published dataset (Rutz, Bisson et al. 2023) from the LOTUS initiative (Rutz, Sorokina et al. 2022), the NPASS database (version NPASS-2023) (Zhao, Yang et al. 2023), and the COCONUT natural product database (accessed 2025-06-09) (Chandrasekhar, Rajan et al. 2025). Additional data were sourced from previous reports. These records were analyzed, and InChI notations were extracted for each *Ferula* natural product.

InChIs were converted to InChI keys and SMILES using RDKit (version 2023.9.5) (Landrum, Tosco et al. 2023). *Ferula* natural products were structurally classified using NPClassifier (Kim, Wang et al. 2021) (, accessed July 13, 2025) with SMILES notations and ClassyFire (Djoumbou Feunang, Eisner et al. 2016) (pyclassyfire API, accessed July 13, 2025) with InChI keys. Metabolites classified into multiple biosynthetic pathways or superclasses were referred to as “hybrids” and categorized under their primary classification. InChI or SMILES notations were used to retrieve chemical properties (CID, molecular formula, IUPAC name, monoisotopic mass, xlogP, and TPSA) from PubChem (accessed July 13, 2025) using PubChemPy (version 1.0.4) (Swain 2014). The Biopython (version 1.82) (Cock, Antao et al. 2009) KEGG Compound package (accessed July 13, 2025) was utilized to annotate InChIs or SMILES notations with KEGG compound IDs, KEGG map IDs, and KEGG map names. PubChem fingerprints were generated for each natural product SMILES using the scikit-fingerprints Python package (version 1.11.0) (Adamczyk and Ludynia 2024).

### 4.6 Cheminformatics, Data Analysis, and Statistics

Tanimoto similarity was calculated using the Python package scikit-fingerprints (version 1.11.0) (Adamczyk and Ludynia 2024). Chemical structures were drawn and annotated using RDKit (version 2023.9.5) (Landrum, Tosco et al. 2023) or the RCSB Chemical Sketch Tool (, accessed January 6, 2025).

Data wrangling, visualization, and statistical analyses were conducted using the R programming language (version 4.3.2) and the “tidyverse” (version 2.0.0) R packages. Principal components analysis (PCA) was performed using the R function prcomp(scaled = TRUE) on the working mass feature table after adding 1 and log2-transforming the data. Permutational multivariate analysis of variance (PERMANOVA) was conducted with the R function adonis2() from the vegan package (version 2.6-8). Venn diagrams were created using the R package VennDiagram (version 1.6.0), and dendrograms were generated with the ggtree package (version 3.10.1) (Yu, Smith et al. 2017). Differential metabolite abundance analysis was performed using the R package limma (version 3.58.1) (Ritchie, Phipson et al. 2015). Differentially abundant metabolites were identified based on a log2-fold change greater than 1 and a false discovery rate (FDR) of less than 0.05.

## 5. Declarations

### Author contributions

Conceptualization, V.D., S.S.; Data curation, S.M., K.S., R.A., and V.D.; Formal analysis, K.A., R.A., SM; Funding acquisition, V.D. and S.S.; Investigation, S.M., K.A, and R.A.; Methodology, V.D., S.M., R.A., and K.A.; Project administration, V.D.; Supervision, V.D.; Validation, K.A., R.A. and V.D.; Visualization, K.A.; Writing— original draft, V.D., S.M; Writing—review and editing, S.S., S.M. V.D., and K.A. All authors have read and approved the final manuscript.

### Data availability

The datasets generated and analyzed during the current study are available in the article and supplementary materials. Complete raw data tables and computational analysis scripts are available at https://doi.org/10.6084/m9.figshare.28585820.v1.

## Acknowledgments

The authors thank the Information Technology Services group at Rutgers University for providing computational resources.

## 6. Short legends for Supporting Information

Short legends for Supporting Information; (x) References; (xi) Tables; (xii) Figure legends; (xiii) Figures.

## References

1. Abaza, S. (2024). “Recent advances in identification of potential drug targets and development of novel drugs in parasitic diseases: Part V: The value of natural products in drug discovery: Helminths.” Parasitologists United Journal 17(2): 57–73.10.21608/puj.2024.292672.1250

2. Adamczyk, J. and P. Ludynia (2024). “Scikit-fingerprints: Easy and efficient computation of molecular fingerprints in Python.” SoftwareX 28: 101944.10.1016/j.softx.2024.101944

3. Ahmad, S. R. and S. Karmakar (2023). “The Role of Medicinal Plants in Drug Discovery across the World.” Ind. J. Pure App. Biosci 11(2): 30–41.10.18782/2582-2845.8995

4. Arumugam, R., B. Elanchezhian, J. Samidurai and K. Amirthaganesan (2022). “Comparative antioxidant, antibacterial and phytochemical analysis of roots, stems, leaves and seeds from *Cleome rutidosperma* DC.” Natural Resources for Human Health 2(4): 479–484.10.53365/nrfhh/146009

5. Attarian, B., G. Kavoosi, Z. Bordbar and H. Sadeghi (2024). “Proximate composition, physico-chemical properties, techno-functional properties, nutritional quality, and functional activity of *Ferula assafoetida* oleo-gum-resin.” Journal of Food Composition and Analysis 129: 106073.10.1016/j.jfca.2024.106073

6. Aydoğan, F., Ş. Baykan and B. D. Bütüner (2019). “Cytotoxic activity of sesquiterpenoids isolated from endemic *Ferula tenuissima* hub.-mor & peşmen.” Turkish Journal of Pharmaceutical Sciences 16(4): 476.10.4274/tjps.galenos.2018.23356

7. Bagheri, S. M. and M. Esmailidehaj (2024). “A Comprehensive Review of the Pharmacological Effects of Genus *Ferula* on Central Nervous System Disorders.” Central Nervous System Agents in Medicinal Chemistry (Formerly Current Medicinal Chemistry-Central Nervous System Agents) 24(2): 105–116.10.2174/0118715249256485231031043722

8. Baimakhanova, B., A. Sadanov, A. Bogoyavlenskiy, V. Berezin, L. Trenozhnikova, G. Baimakhanova, A. Ibraimov, E. Serikbayeva, Z. Arystanov and T. Arystanova (2025). “Exploring phytochemicals and their pharmacological applications from ethnomedicinal plants: A focus on Lycium barbarum, Solanacea.” Heliyon.10.1016/j.heliyon.2025.e41782

9. Baykan, S., F. Aydogan, M. Ozturk, B. Debelec-Butuner, Ç. Yengin and B. Ozturk (2020). “Ferutinin content and cytotoxic effects of various *Ferula* L. species on prostate cancer (PC-3) cell line.” J. Res. Pharm 24: 142–149.10.3533/jrp.2020.120

10. Bellavite, P. (2023). “Neuroprotective potentials of flavonoids: Experimental studies and mechanisms of action.” Antioxidants 12(2): 280.10.3390/antiox12020280

11. Bogoyavlenskiy, A., P. Alexyuk, M. Alexyuk, V. Berezin, I. Zaitseva, E. Omirtaeva, A. Manakbayeva, Y. Moldakhanov, E. Anarkulova and A. Imangazy (2024). “The Ability of Combined Flavonol and Trihydroxyorganic Acid to Suppress SARS-CoV-2 Reproduction.” Viruses 17(1): 37.10.3390/v17010037

12. Chandrasekhar, V., K. Rajan, S. R. S. Kanakam, N. Sharma, V. Weißenborn, J. Schaub and C. Steinbeck (2025). “COCONUT 2.0: a comprehensive overhaul and curation of the collection of open natural products database.” Nucleic Acids Research 53(D1): D634–D643.10.1093/nar/gkae1063

13. Ciumărnean, L., M. V. Milaciu, O. Runcan, Ș. C. Vesa, A. L. Răchișan, V. Negrean, M.-G. Perné, V. I. Donca, T.-G. Alexescu and I. Para (2020). “The effects of flavonoids in cardiovascular diseases.” Molecules 25(18): 4320.10.3390/molecules25184320

14. Cock, P. J., T. Antao, J. T. Chang, B. A. Chapman, C. J. Cox, A. Dalke, I. Friedberg, T. Hamelryck, F. Kauff and B. Wilczynski (2009). “Biopython: freely available Python tools for computational molecular biology and bioinformatics.” Bioinformatics 25(11): 1422.10.1093/bioinformatics/btp163

15. Das, A. and R. Ruhal (2025). “Potential of plants-based alkaloids, terpenoids and flavonoids as antibacterial agents: An update.” Process Biochemistry.10.1016/j.procbio.2025.01.003

16. Das, D., N. Keerthi, A. Banerjee, A. K. Duttaroy and S. Pathak (2025). “An Overview of Cardiovascular Disease Management with Plant-Derived Bioactive Compounds.” Plant Derived Bioactive Compounds in Human Health and Disease: 156–176

17. De Luca, V., V. Salim, S. M. Atsumi and F. Yu (2012). “Mining the biodiversity of plants: a revolution in the making.” Science 336(6089): 1658–1661.10.1126/science.1217410

18. Djoumbou Feunang, Y., R. Eisner, C. Knox, L. Chepelev, J. Hastings, G. Owen, E. Fahy, C. Steinbeck, S. Subramanian and E. Bolton (2016). “ClassyFire: automated chemical classification with a comprehensive, computable taxonomy.” Journal of cheminformatics 8: 1–20.10.1186/s13321-016-0174-y

19. Dührkop, K., M. Fleischauer, M. Ludwig, A. A. Aksenov, A. V. Melnik, M. Meusel, P. C. Dorrestein, J. Rousu and S. Böcker (2019). “SIRIUS 4: a rapid tool for turning tandem mass spectra into metabolite structure information.” Nature methods 16(4): 299–302.10.1038/s41592-019-0344-8

20. Dushenkov, V. and A. Dushenkov (2022). “Botanicals as prospective agents against SARS-CoV-2 virus.” Avicenna Bulletin (Vestnik Avitsenny) 24(1): 113–122.10.25005/2074-0581-2022-24-1-113-122

21. Elarabany, N., A. Hamad and N. M. Alzamel (2023). “Antitumor and Phytochemical Properties of *Ferula assa-foetida* L. Oleo-Gum–Resin against HT-29 Colorectal Cancer Cells In Vitro and in a Xenograft Mouse Model.” Molecules 28(24): 8012.10.3390/molecules28248012

22. Flores-Morales, V., A. P. Villasana-Ruíz, I. Garza-Veloz, S. González-Delgado and M. L. Martinez-Fierro (2023). “Therapeutic effects of coumarins with different substitution patterns.” Molecules 28(5): 2413.10.3390/molecules28052413

23. Ghaforzoda, L., F. Sharopov, I. Gulmurodov, M. Bakri, S. Numonov and H. Aisa (2025). “Volatile Metabolites in Essential Oil of *Ferula violacea*.” Chemistry of Natural Compounds: 1–3. 10.1007/s10600-025-04609-2

24. Hart, C. E., Y. Gadiya, T. Kind, C. A. Krettler, M. Gaetz, B. B. Misra, D. Healey, A. Allen, V. Colluru and D. Domingo-Fernández (2025). “Defining the limits of plant chemical space: challenges and estimations.” GigaScience 14: giaf033.10.1093/gigascience/giaf033/8106437

25. Hassanein, E. H., H. S. Althagafy, M. A. Baraka, E. K. Abd-Alhameed, I. M. Ibrahim, M. S. Abd El-Maksoud, N. M. Mohamed and S. A. Ross (2024). “The promising antioxidant effects of lignans: Nrf2 activation comes into view.” Naunyn-Schmiedeberg’s Archives of Pharmacology 397(9): 6439–6458.10.1007/s00210-024-03102-x

26. Heinrich, M., J. Mah and V. Amirkia (2021). “Alkaloids used as medicines: Structural phytochemistry meets biodiversity—An update and forward look.” Molecules 26(7): 1836.10.3390/molecules26071836

27. Hoffmann, M. A., L.-F. Nothias, M. Ludwig, M. Fleischauer, E. C. Gentry, M. Witting, P. C. Dorrestein, K. Dührkop and S. Böcker (2022). “High-confidence structural annotation of metabolites absent from spectral libraries.” Nature Biotechnology 40(3): 411–421.10.1038/s41587-021-01045-9

28. Ikeda, K., Y. Arao, H. Otsuka, S. Nomoto, H. Horiguchi, S. Kato and F. Kayama (2002). “Terpenoids found in the Umbelliferae family act as agonists/antagonists for ERα and ERβ: Differential transcription activity between ferutinine-liganded ERα and ERβ.” Biochemical and biophysical research communications 291(2): 354–360.10.1006/bbrc.2002.6446

29. Jiang, M., M. Peng, Y. Li, G. Li, X. Li and L. Zhuang (2024). “Quality evaluation of four *Ferula* plants and identification of their key volatiles based on non-targeted metabolomics.” Frontiers in Plant Science 14: 1297449.10.3389/fpls.2023.1297449

30. Karimi, M. R., P. Jariani, J.-L. Yang and M. R. Naghavi (2024). “A comprehensive review of the molecular and genetic mechanisms underlying gum and resin synthesis in *Ferula* species.” International Journal of Biological Macromolecules: 132168.10.1016/j.ijbiomac.2024.132168

31. Khosnutdinova, T., N. Gemejiyeva, Z. Z. Karzhaubekova and N. Sultanova (2023). “Coumarins of genus *Ferula* L.(Apiaceae Lindl.).” Eurasian Chemico-Technological Journal 25(1): 39–56.10.18321/ectj1494

32. Kim, H. W., M. Wang, C. A. Leber, L.-F. Nothias, R. Reher, K. B. Kang, J. J. Van Der Hooft, P. C. Dorrestein, W. H. Gerwick and G. W. Cottrell (2021). “NPClassifier: a deep neural network-based structural classification tool for natural products.” Journal of Natural Products 84(11): 2795–2807.10.1021/acs.jnatprod.1c00399

33. Kir’yanova, I., Y. E. Sklyar, M. Pimenov and Y. V. Baranova (1979). “Terpenoid coumarins of *Ferula violacea* and *F. eugenii*.” Chemistry of Natural Compounds 15(4): 499–499

34. Landrum, G., P. Tosco, B. Kelley, Ric, D. Cosgrove, Sriniker, R. Vianello, Gedeck, N. Schneider, G. Jones, E. Kawashima, D. N. A. Dalke, B. Cole, M. Swain, S. Turk, A. Savelev, A. Vaucher, M. Wójcikowski, I. Take, V. F. Scalfani, D. Probst, K. Ujihara, g. Godin, A. Pahl, R. Walker, J. Lehtivarjo, F. Berenger, Strets and D. Jason (2023). rdkit/rdkit: Release_2023.09.5.10.5281/zenodo.10633624

35. Letchuman, S., H. D. Madhuranga, M. Kaushalya, A. D. Premarathna and M. Saravanan (2024). “Alkaloids Unveiled: A Comprehensive Analysis of Novel Therapeutic Properties, Mechanisms, and Plant-Based Innovations.” Intelligent Pharmacy.10.1016/j.ipha.2024.09.007

36. Li, J., X. Ma, L. Luo, D. Tang and L. Zhang (2023). “The What and Who of Dietary Lignans in Human Health: Special Attention to Estrogen Effects and Safety Evaluation.” Journal of Agricultural and Food Chemistry 71(44): 16419–16434.10.1021/acs.jafc.3c02680J

37. Mishra, S. (2024). “Neuroprotective Effect of Naturally Occurring Flavonoids.” Central Nervous System Agents in Medicinal Chemistry.10.2174/0118715249344284241112184703

38. Mohammadi, R., H. Forouzanfar, H. Rahimi, S.-M. Mohamadi-Zarch, K. Jamhiri and S. M. Bagheri (2025). “Antiviral Effect of *Ferula* Plants and their Potential for Treatment of COVID-19: A Comprehensive Review.” Current pharmaceutical biotechnology.10.2174/0113892010285343240530040218

39. Nouioura, G., M. El Fadili, A. El Barnossi, E. H. Loukili, H. Laaroussi, M. Bouhrim, J. P. Giesy, M. A. Aboul-Soud, Y. A. Al-Sheikh and B. Lyoussi (2024). “Comprehensive analysis of different solvent extracts of *Ferula communis* L. fruit reveals phenolic compounds and their biological properties via in vitro and in silico assays.” Scientific Reports 14(1): 8325.10.1038/s41598-024-59087-3

40. Nowak, A., S. Świerszcz, S. Nowak, H. Hisorev, E. Klichowska, A. Wróbel, A. Nobis and M. Nobis (2020). “Red List of vascular plants of Tajikistan–the core area of the Mountains of Central Asia global biodiversity hotspot.” Scientific Reports 10(1): 6235.10.1038/s41598-020-63333-9

41. Pimenov, M. G. (2020). “Updated checklist of the Umbelliferae of Middle Asia and Kazakhstan: nomenclature, synonymy, typification, distribution.” Turczaninowia 23(4): 127–257.10.14258/turczaninowia.23.4.12

42. Reaxys, E. (2024). “Reaxys chemical database.” Retrieved October 3, 2024, 2024, from https://www.elsevier.com/products/reaxys.

43. Ritchie, M. E., B. Phipson, D. Wu, Y. Hu, C. W. Law, W. Shi and G. K. Smyth (2015). “limma powers differential expression analyses for RNA-sequencing and microarray studies.” Nucleic acids research 43(7): e47–e47.10.1093/nar/gkv007

44. Rutz, A., J. Bisson and P.-M. Allard (2023). The LOTUS Initiative for Open Natural Products Research: frozen dataset union wikidata (with metadata).10.5281/zenodo.7534071

45. Rutz, A., M. Sorokina, J. Galgonek, D. Mietchen, E. Willighagen, A. Gaudry, J. G. Graham, R. Stephan, R. Page and J. Vondrášek (2022). “The LOTUS initiative for open knowledge management in natural products research.” Elife 11: e70780.10.7554/eLife.70780

46. Safarov, N. M., H. H. Hisoriev and K. Shermatov (2019). “Plant diversity in Tajikistan and present conservation status of their rare and endangered species.” Известия Академии наук Республики Таджикистан. Отделение биологических и медицинских наук(4): 7–20

47. Sahoo, M. R., R. Raghava Varrier, B. S. Harihara Sharma, A. Rajendran and B. Guru (2024). “Phytochemical Profiling for Leaf, Stem, Roots, and Seeds of Withania somnifera (L.) Dunal.” J Rep Pharm Sci 12(1): e146161.10.5812/jrps.146161

48. Saidkhodzhaev, A. I. and G. K. Nikonov (1973). “The structure of ferutinin.” Chemistry of Natural Compounds 9(1): 25–26.10.1007/BF00580882

49. Satorov, S., S. Mavlonazarova, A. P. Bogoyavlenskiy, S. D. Yusufi and V. Dushenkov (2024). “Efficacy of *Ferula* L. species extracts from Tajikistan against influenza viruses.” Afr. J. B.Sc. 6(9): 3254–3268.10.33472/AFJBS.6.9.2024.3217-3245

50. Satorov, S., S. Mavlonazarova, S. Yusufi and V. Dushenkov (2024). “Total Polyphenol Content, Antioxidant Potential, Antibacterial and Antifungal Properties of *Ferula* L. Species Growing in Tajikistan.” Journal of Drug and Alcohol Research 13(9).10.4303/JDAR/236424

51. Schläger, S. and B. Dräger (2016). “Exploiting plant alkaloids.” Current opinion in biotechnology 37: 155–164.10.1016/j.copbio.2015.12.003

52. Schrimpe-Rutledge, A. C., S. G. Codreanu, S. D. Sherrod and J. A. McLean (2016). “Untargeted metabolomics strategies—challenges and emerging directions.” Journal of the American Society for Mass Spectrometry 27(12): 1897–1905.10.1007/s13361-016-1469-y

53. Schymanski, E. L., J. Jeon, R. Gulde, K. Fenner, M. Ruff, H. P. Singer and J. Hollender (2014). Identifying small molecules via high resolution mass spectrometry: communicating confidence, ACS Publications. dx.doi.org/10.1021/es5002105 | Environ. Sci. Technol. 2014, 48, 2097−2098

54. Seregin, A. P. E. (2023). “Moscow Digital Herbarium: Electronic resource.” Retrieved 03/02/2023, 2024, from https://plant.depo.msu.ru/.

55. Sharopov, F. and W. N. Setzer (2018). “Medicinal plants of Tajikistan.” Vegetation of Central Asia and environs: 163–209

56. Shende, V. V., K. D. Bauman and B. S. Moore (2024). “The shikimate pathway: gateway to metabolic diversity.” Natural Product Reports 41(4): 604–648.10.1039/d3np00037k

57. Swain, M. (2014). “PubChemPy.” Retrieved December 23, 2024, from https://github.com/mcs07/PubChemPy.

58. Wang, J., Q. Zheng, H. Wang, L. Shi, G. Wang, Y. Zhao, C. Fan and J. Si (2023). “Sesquiterpenes and Sesquiterpene Derivatives from *Ferula*: Their Chemical Structures, Biosynthetic Pathways, and Biological Properties.” Antioxidants 13(1): 7 10.3390/antiox13010007

59. WFO. (2024). “World Flora Online.” Retrieved 25 July, 2024, from https://wfoplantlist.org.

60. Yadav, N. K., A. B. Patel, S. Baidya and P. Biswas (2025). “Interactive effects of solvent extraction and drying methods on LC-ESI-QTOF-MS/MS-based characterization of bioactive phenolics and antioxidant activities in duckweed (Wolffia globosa).” European Food Research and Technology: 1–20.10.1007/s00217-024-04647-0

61. Younes, A. H. and Y. F. Mustafa (2024). “Plant - Derived Coumarins: A Narrative Review of Their Structural and Biomedical Diversity.” Chemistry & Biodiversity 21(6): e202400344.10.1002/cbdv.202400344

62. Yu, G., D. K. Smith, H. Zhu, Y. Guan and T. T. Y. Lam (2017). “GGTREE: an R package for visualization and annotation of phylogenetic trees with their covariates and other associated data.” Methods in Ecology and Evolution 8(1): 28–36.10.1111/2041-210X.12628

63. Zhao, H., Y. Yang, S. Wang, X. Yang, K. Zhou, C. Xu, X. Zhang, J. Fan, D. Hou and X. Li (2023). “NPASS database update 2023: quantitative natural product activity and species source database for biomedical research.” Nucleic Acids Research 51(D1): D621–D628.10.1093/nar/gkac1069

64. Zheng, X., X. Zhang and F. Zeng (2025). “Biological Functions and Health Benefits of Flavonoids in Fruits and Vegetables: A Contemporary Review.” Foods 14(2): 155.10.3390/foods14020155

